# IRF2BP2 counteracts the ATF7/JDP2 AP-1 heterodimer to prevent inflammatory overactivation in acute myeloid leukemia (AML) cells

**DOI:** 10.1101/2023.06.09.544165

**Authors:** Sabrina Fischer, Lisa M. Weber, Bastian Stielow, Miriam Frech, Clara Simon, Julie Könnecke, Ignasi Forné, Andrea Nist, Uta Maria Bauer, Thorsten Stiewe, Andreas Neubauer, Robert Liefke

**Affiliations:** Institute of Molecular Biology and Tumor Research (IMT), Philipps University of Marburg, Marburg 35043, Germany; Department of Hematology, Oncology, and Immunology, University Hospital Giessen and Marburg, Marburg 35043, Germany; Protein Analysis Unit, Biomedical Center (BMC), Faculty of Medicine, Ludwig-Maximilians-University (LMU) Munich, Martinsried 82152, Germany; Genomics Core Facility, Institute of Molecular Oncology, Member of the German Center for Lung Research (DZL), Philipps University of Marburg, Marburg 35043, Germany

**Keywords:** acute myeloid leukemia, IRF2BP2, inflammation, interferon, ATF7, JDP2, transcription, AP-1

## Abstract

Acute myeloid leukemia (AML) is a hematological malignancy characterized by abnormal proliferation and accumulation of immature myeloid cells in the bone marrow. Inflammation plays a crucial role in AML progression, but excessive activation of cell-intrinsic inflammatory pathways can also trigger cell death. IRF2BP2 is a chromatin regulator implicated in AML pathogenesis, although its precise role in this disease is not fully understood. In this study, we demonstrate that IRF2BP2 interacts with the AP-1 heterodimer ATF7/JDP2, which is involved in activating inflammatory pathways in AML cells. We show that IRF2BP2 is recruited by the ATF7/JDP2 dimer to chromatin and counteracts its gene-activating function. Loss of IRF2BP2 leads to overactivation of inflammatory pathways, resulting in strongly reduced proliferation. Our research indicates that a precise equilibrium between activating and repressive transcriptional mechanisms creates a pro-oncogenic inflammatory environment in AML cells. The ATF7/JDP2-IRF2BP2 regulatory axis is likely a key regulator of this process and may therefore represent a promising therapeutic vulnerability for AML. Thus, our study provides new insights into the molecular mechanisms underlying AML pathogenesis and identifies a potential therapeutic target for AML treatment.

## Introduction

Acute myeloid leukemia (AML) is a devastating disease that predominantly affects adults and progresses rapidly, often leading to death within weeks or months if left untreated^1^. The disease is characterized by the hyperproliferation of undifferentiated myeloid progenitor cells caused by genetic alterations that affect cytoplasmic tyrosine kinases, signaling modulators, and epigenetic regulators^1, 2^. While progress has been made in identifying and targeting specific cellular pathways affected by genetic alterations, the survival of AML patients remains poor, with frequent relapse after the first remission and the development of drug resistance^1, 3^. Therefore, new therapeutic approaches are urgently needed to improve the outcome for AML patients.

Inflammation is a critical factor in the pathogenesis of AML and has become an attractive therapeutic target in recent years^4^. IRF2BP2, a protein that regulates inflammation pathways, has emerged as an important player in AML^5^. Mutation of IRF2BP2 has also been implicated in immunodeficiency disorders^6^. Originally, IRF2BP2 was identified together with its homolog IRF2BP1, as a transcriptional corepressor for the interferon regulatory factor 2 (IRF2), thus the naming of the protein^7^. An additional homolog is called IRF2BPL (also known as EAP1). These three proteins are commonly found together^8, 9^ and exert a repressive function in gene regulation^10^. Biochemically, all three proteins are characterized by the presence of an N-terminal zinc finger and a C-terminal RING domain. The zinc finger has been demonstrated to be involved in interconnecting the three proteins, allowing the formation of larger protein complexes^8^. The RING domain has been implicated in ubiquitin ligase activity^11, 12^. The repressive role of IRF2BP2 has been shown to be independent of histone deacetylases^7^. Instead, E3 ubiquitin ligase activity as well as sumoylation mechanisms have been implicated in the repressive activity of IRF2BP2^12, 13^. Besides IRF2, the IRF2BP2 complex interacts with a variety of other transcription factors and chromatin regulators^10^, including KLF2^14^, NFAT1^15^, ETO2^9^, HNF4α^12^, and TEAD4^16^. However, the specific transcription factors involved in recruiting IRF2BP2 to chromatin in AML cells and how they may work together to establish a pro-proliferative transcriptional setting in AML remain undetermined.

Here, we provide further evidence of the crucial role of IRF2BP2 in AML by demonstrating its specific interaction with the AP-1 heterodimer, composed of ATF7 and JDP2. We show that IRF2BP2 localizes with ATF7 genome-wide and that its efficient chromatin binding depends on ATF7 and JDP2. ATF7/JDP2 activates inflammatory genes in AML cells, and IRF2BP2 represses this transcriptional activity. When IRF2BP2 is absent, overactivation of inflammatory pathways occurs, inhibiting cellular proliferation and eventually leading to cell death. Our findings suggest that targeting the ATF7/JDP2-IRF2BP2-axis may be a promising therapeutic approach by modulating inflammation and promoting cell death in leukemic cells.

## Results

### IRF2BP2 is essential for the growth of acute myeloid leukemia cells

Using publicly available CRISPR perturbation data^17^, we previously identified novel proteins essential for AML cell proliferation, including IRF8^18^. Upon this analysis, we also identified the protein IRF2BP2 (Interferon regulatory factor 2 binding protein 2) as a potentially critical factor in AML. In the CRISPR screen, IRF2BP2 scores strongly negative in AML cell lines, but not in other human cell lines (**Fig. 1a**), suggesting that deletion of IRF2BP2 impairs the proliferation of AML cells. Furthermore, IRF2BP2 is commonly upregulated in AML (**Fig. 1b**) and can easily be detected in AML cell lines (**Supplementary Figure 1a,b**). Investigation of public patient survival data indicates that high IRF2BP2 expression is typically associated with a poorer prognosis (**Fig. 1c, d**).

**Fig. 1.**
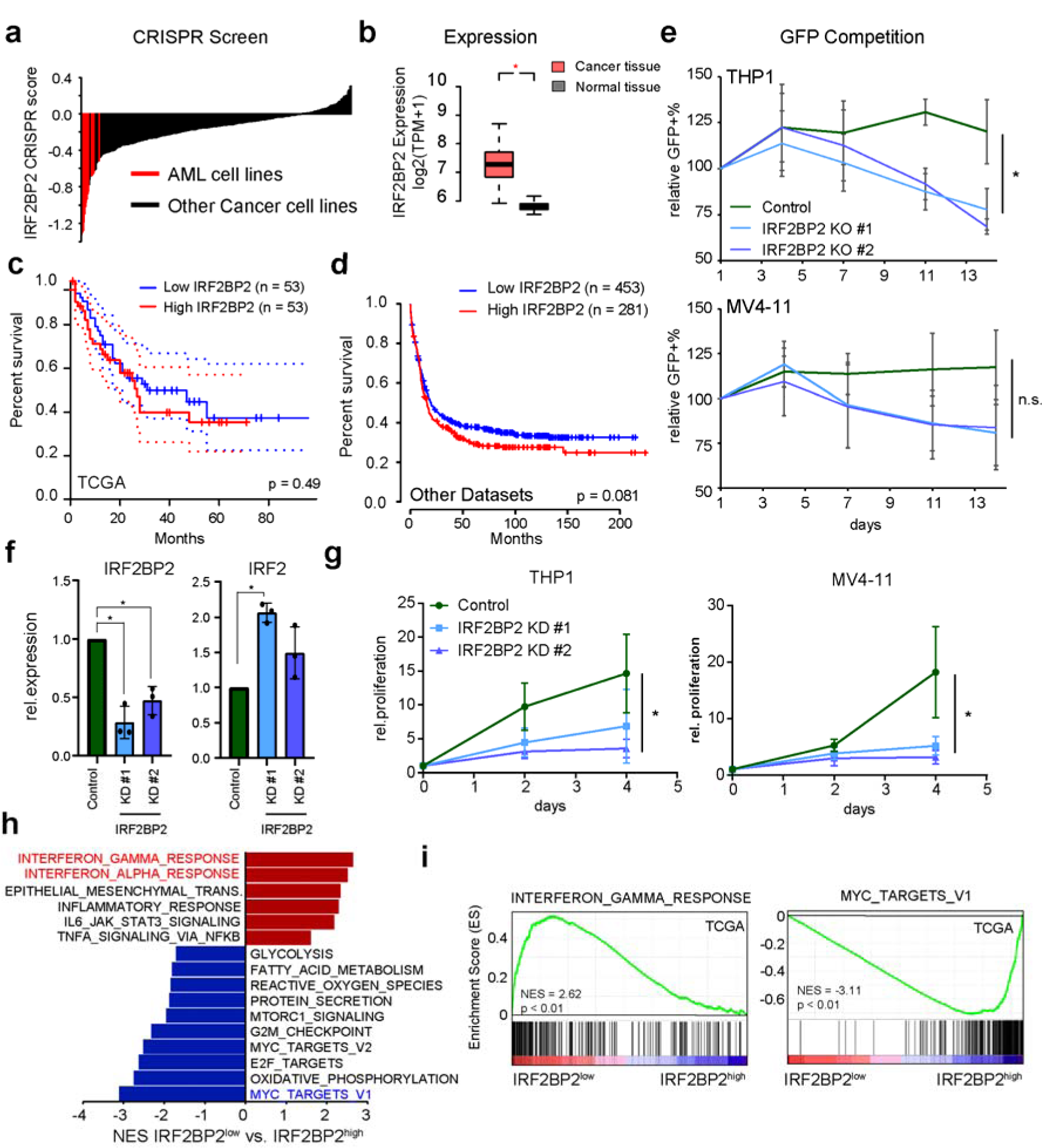
IRF2BP2 is required for AML cell proliferation. a) Distribution of CRISPR scores upon IRF2BP2 deletion in 357 human cancer cell lines^17^, with AML cell lines marked in red. b) Expression of IRF2BP2 in AML versus non-AML samples, using data from GePIA^21^. c) Kaplan-Meier survival curve (disease-free survival) of AML patients dependent on IRF2BP2 expression. Data were derived from TCGA^19^ and were visualized using GePIA^21^. d) Kaplan-Meier survival curve (overall survival) of AML patients dependent on IRF2BP2 expression. The data base on multiple microarray datasets, visualized by the Kaplan-Meier-Plotter^22^ using “auto select best cutoff”. e) Negative-selection competition assay showing the percentage of GFP+ control- or sgKAT6A-transduced cells over time (n=2–3 each group). Data represent the mean +/− s.d. of at least two biological replicates. P-values were evaluated via ANOVA. f) Expression of IRF2BP2 and IRF2 after IRF2BP2 shRNA-mediated knockdown. Data represent the mean +/− s.d. of three biological replicates. P-values were evaluated via ANOVA. g) Growth of THP1 and MV4-11 cells after knockdown of IRF2BP2. Data represent the mean +/− s.d. of at least three biological replicates. h) GSEA comparing AML samples from TCGA^19^, with low versus high IRF2BP2 expression. i) GSEA plots of the two top up- and downregulated pathways from h).

To investigate the role of IRF2BP2 in AML cells in more detail, we first performed CRISPR/Cas9-mediated deletion of IRF2BP2 in several AML cell lines, such as THP1 and MV4-11. Consistent with the results from the CRISPR-Screen, we observed cell death and failed to obtain stable KO cells. To confirm the impact of IRF2BP2 knockout, we investigated the survival of knockout cells, additionally expressing GFP as a marker relative to GFP-negative cells. Deletion of IRF2BP2 led to a loss of the GFP-positive cell population (**Fig. 1e**), suggesting that the removal of IRF2BP2 impairs cell growth. To circumvent the death after knockout, we used an shRNA-mediated knockdown approach, which reduced the IRF2BP2 mRNA level to approximately 30% of the wild-type level (**Fig. 1f**). In these cells, we observed an upregulation of IRF2, supporting an interplay between IRF2BP2 and IRF2-mediated interferon pathways. Knockdown led to an impaired proliferation of the cells (**Fig. 1g**), demonstrating that high IRF2BP2 levels are important to sustain high proliferation of the investigated AML cells and that low levels of IRF2BP2 prevent optimal growth.

To gain insights into the underlying reason, we made use of AML patient data from TCGA^19^. We found that low IRF2BP2 expression correlated with increased expression of genes related to the interferon response and inflammation (**Fig. 1h, i**), consistent with the elevated level of IRF2 upon knockdown (**Fig. 1f**). Moreover, low IRF2BP2 is linked to decreased expression of MYC target genes and metabolic pathways, typically important for rapid cancer cell growth^20^(**Fig. 1h,j**). These findings support that high IRF2BP2 expression is involved in setting up a pro-proliferative transcriptional network in AML cells and that reducing the level of IRF2BP2 enhances inflammatory pathways, which may contribute to cell death. However, the molecular processes by which IRF2BP2 regulates this cellular program in AML cells and why its presence is so crucial for survival are not fully understood.

### IRF2BP2-complex interacts with the ATF7/JDP2 dimer in AML cells

IRF2BP2 is known to interact with various transcription factors^8, 9, 12, 14^, but with which factors it interacts in the context of AML is unknown. To identify interaction partners of IRF2BP2 in AML cells, we expressed Flag-HA-tagged IRF2BP2 (Isoform B, UniProt: Q7Z5L9-2) in THP1 and MV4-11 AML cells, where IRF2BP2 is found in the nucleus and at the chromatin (**Supplementary Fig. 1b, c**), and purified IRF2BP2 using tandem-affinity purification. Flag-HA-tagged GFP served as a negative control. IRF2BP2 forms a stable complex and colocalizes with its partner IRF2BP1 in both cell lines (**Fig. 2a**, **Supplementary Fig. d**), demonstrating the validity of this approach. Additionally, we recapitulated previous reports that IRF2BP2 interacts with itself^8^ (**Fig. 2a**).

**Fig. 2:**
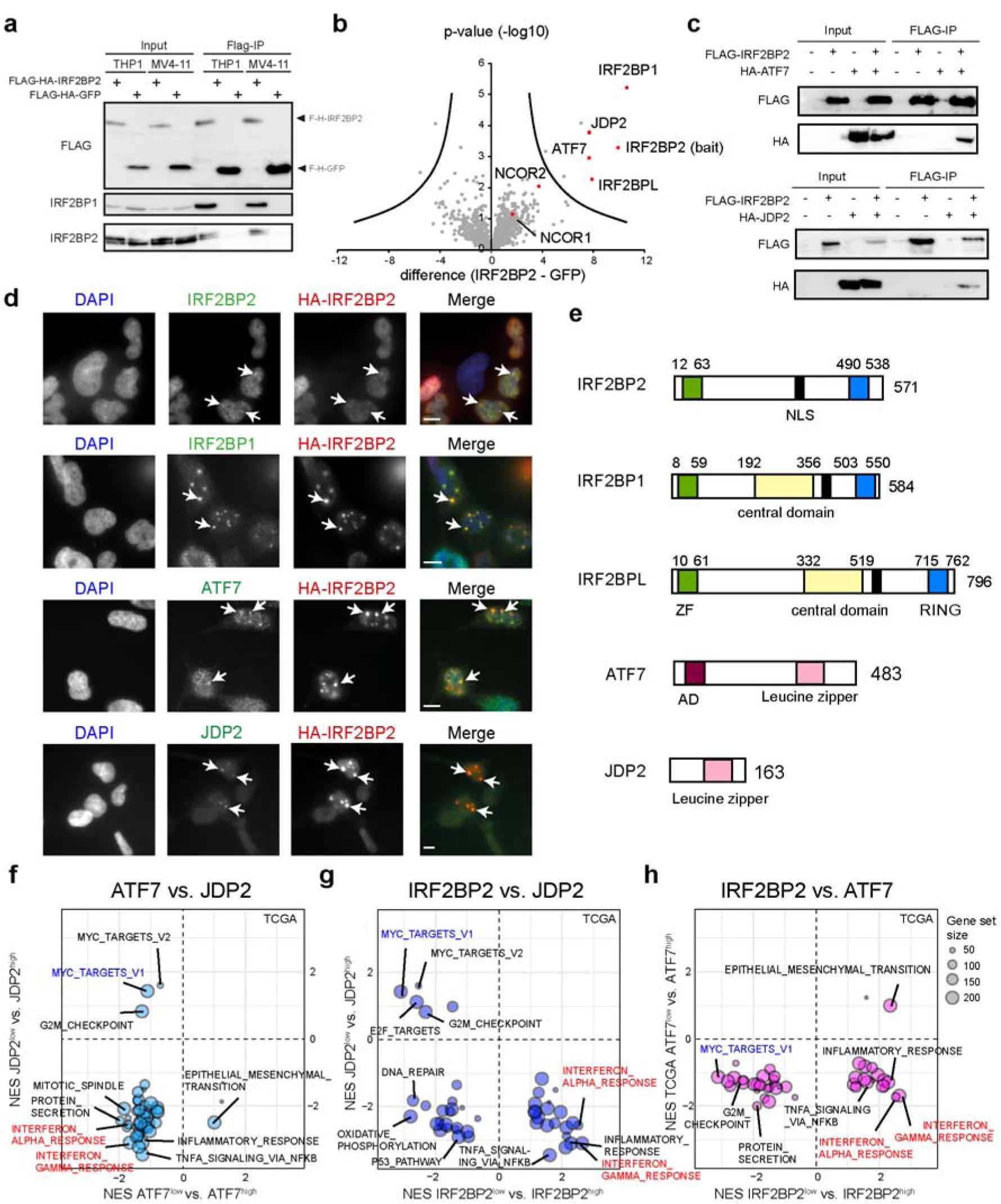
The ATF7/JDP2 dimer interacts with IRF2BP2 but has an opposing role in regulating inflammatory pathways. a) Semi-endogenous co-immunoprecipitation showing the interaction of IRF2BP2 with itself and IRF2BP1. b) Semi-quantitative mass-spectrometry after co-immunoprecipitation of Flag-IRF2BP2 in THP1 cells. c) Confirmation of interaction of IRF2BP2 with ATF7 and JDP2 using co-immunoprecipitation experiments in HEK293 cells. d) Immunofluorescence showing the colocalization of ectopically expressed IRF2BP2 with IRF2BP1, JDP2 and ATF7 in nuclear foci (white arrow) in HEK293 cells. e) Domain structure of IRF2BP2, IRF2BP1, IRF2BPL, ATF7 and JDP2. See also Supplementary Fig. S2. NLS = Nuclear localization signal, ZF = zinc finger domain, RING = RING finger domain f) Bubble plot showing the gene sets (hallmarks) correlating with high ATF7 and JDP2 in TCGA AML samples. Inflammatory pathways negatively correlate with low ATF7 and JDP2 expression (lower left quadrant). g) Bubble plot showing the correlation of gene sets that correlate with low JDP2 and IRF2BP2 expression in TCGA samples. Inflammatory pathways anti-correlate (lower right quadrant). h) As in g), but the comparison between low ATF7 and low IRF2BP2 expression is shown. Inflammatory pathways anti-correlate (lower right quadrant).

Subsequently, we investigated IRF2BP2-associated proteins in THP1 cells using label-free quantitative mass spectrometry (**Fig. 2b**). In this experiment, IRF2BP1 and IRF2BPL were most strongly enriched, suggesting that they are the main interactors of IRF2BP2, consistent with previous reports^8^. Via co-immunoprecipitation experiments, we validated that both IRF2BP1 and IRF2BP2 interact with IRF2BPL (**Supplementary Fig. 1e**). In the following, we refer to the “IRF2BP2 complex” when talking about these three proteins together. Looking at the other enriched proteins, we did not find IRF2 or other described interaction partners, except for NCOR1 and NCOR2^9^. Instead, we identified the transcription factors ATF7 and JDP2 to be highly enriched (**Fig. 2b**). ATF7 and JDP2 belong to the AP-1 transcription factor family and typically act as dimers on DNA^23^. Given that we did not identify further members of the AP-1 family, it suggests that IRF2BP2 specifically interacts with the ATF7/JDP2 dimer but not with other AP-1 dimers.

Using co-immunoprecipitation experiments, we confirmed that IRF2BP2, IRF2BP1, and IRF2BPL can interact with ATF7 and JDP2 (**Fig. 2c, Supplementary Fig. 1f, g**). Notably, during these experiments, we consistently observed that overexpression of JDP2, but not ATF7, led to a strong downregulation of IRF2BP1, IRF2BP2 and IRF2BPL at the protein level (**Fig. 2c**, **Supplementary Fig. 1g,h**). This unexpected effect partially impaired the interpretation of the co-immunoprecipitation experiments, but it further supports that JDP2 is an interactor of these three proteins. However, the reason why JDP2 expression negatively affects the protein level of IRF2BP2 complex members is currently unclear, given that JDP2 has, besides its leucine zipper, no other obvious functional domain (**Fig. 2e**). Thus, the source of this effect requires further investigation. In addition, we observed changes in *IRF2BP1*, *JDP2,* and *IRF2BPL* expression levels upon IRF2BP2 knockdown (**Supplementary Fig. 1i**), raising the possibility that feedback mechanisms are in place to regulate the proper balance of these proteins.

That IRF2BP2 can associate with IRF2BP1, ATF7, and JPD2 is further supported by the observation that ectopically expressed IRF2BP2 colocalizes with the endogenous IRF2BP1, ATF7, and JDP2 in nuclear foci, as demonstrated by immunofluorescence (**Fig. 2d**).

Upon inspection of all identified proteins using the AlphaFold database^24, 25^, we found that IRF2BP1 and IRF2BPL possess a currently unannotated domain in the center of the protein (**Fig. 2e**, **Supplementary Fig. 2a, b**). Furthermore, AlphaFold2^25, 26^ predicts with high confidence that these domains can homo- and heterodimerize, via an intertwined interaction (**Supplementary Fig. S2c, d**). We confirmed this prediction experimentally using co-immunoprecipitation experiments (**Supplementary Fig. 2e**). Additionally, we could demonstrate that dimerization of IRF2BP2, which does not possess the central domain, requires the zinc finger domain, as previously described (**Supplementary Fig. 2f**)^8^. In contrast, IRF2BP1 and IRF2BPL can also interact with themselves without the zinc finger domains (**Supplementary Fig. 2g, h**), suggesting that they can self-associate via both the zinc finger domain and the central domain. This suggests that these proteins exhibit a dual mechanism of self-interaction, utilizing both the zinc finger domain and the central domain. Furthermore, we found that deletion of the central domain reduces the chromatin binding of IRF2BP1 and that this domain alone is present in the chromatin fraction (**Supplementary Fig. 2i**). Thus, this feature may be important for the formation of larger IRF2BP-containing multi-protein complexes and chromatin binding.

### Low ATF7 and JDP2 expression correlates with poorer prognosis and diminished inflammatory pathways in AML

To assess the relevance of the interaction partners of IRF2BP2 in the context of AML, we investigated publicly available data, as before for IRF2BP2 (**Fig. 1**). We found that next to IRF2BP2, also IRF2BP1, ATF7, and JDP2 have predominantly a negative CRISPR score in AML cell lines (**Supplementary Fig. 3a**), indicating that they are also important for proper AML cell proliferation. Only IRF2BPL scores predominantly positive in AML cell lines. Looking at gene expression, we found that both ATF7 and JDP2 are more highly expressed in AML cells than non-cancerous control, while IRF2BP1 and IRF2BPL are not particularly highly expressed in AML samples (**Supplementary Fig. 3b**). Notably, the gene expression levels of these proteins are all lower compared to IRF2BP2 (compare **Fig. 1b** and **Supplementary Fig. 3b**), which may explain why IRF2BP2 deletion has a particularly strong impact on cell proliferation, as seen by the more negative CRISPR scores for IRF2BP2 (compare **Fig. 1a** and **Supplementary Fig. 3a**). Nonetheless, high expression of ATF7, JDP2, and IRF2BPL correlates with a poorer prognosis, as well, while IRF2BP1 expression is not prognostic (**Supplementary Fig. 3c**). In sum, this analysis supports that the identified interaction partners of IRF2BP2 are all expressed in AML cells and that their high expression has a predominantly positive influence on cell proliferation and a negative impact on patient survival.

To better understand the underlying role of ATF7 and JDP2 in AML cells, we investigated AML gene expression data from TCGA^19^. We compared AML samples with low versus high ATF7 and JDP2 expression, respectively, using GSEA. We found that their low expression correlates with low expression of genes involved in interferon pathways and inflammation (lower left quadrant) (**Fig. 2f**). This result is opposite to what we previously found IRF2BP2 (**Fig. 1h, i**), where low expression is linked to higher expression of interferon pathway genes. Consistently, the interferon gene sets anti-correlate between IRF2BP2 and ATF7 and JDP2, respectively (**Fig. 2g, h**, lower right quadrants). Thus, these results suggest that although IRF2BP2 interacts with ATF7 and JDP2, they have opposing roles in regulating inflammatory pathways in AML.

### The IRF2BP2 complex associates with the ATF7-JDP2 dimer likely via multiple interactions

To address the molecular basis of how the IRF2BP2 complex interacts with the ATF7/JDP2 dimer, we performed co-immunoprecipitation experiments in HEK293 cells. In addition to IRF2BP2, we included IRF2BP1, the most enriched other member of the IRF2BP2 complex in our mass-spectrometry experiment (**Fig. 2b**). We confirmed that IRF2BP2 interacts with IRF2BP1, and vice versa, via the N-terminal zinc-finger domain (**Fig. 3a,b**). This is consistent with previous reports, showing that the zinc finger domains are involved in hetero- and homodimerization^8^. Deleting the central domain of IRF2BP1 did not impair the interaction with IRF2BP2 (**Fig. 3b**), suggesting that this domain is irrelevant for the interaction between IRF2BP2 and IRF2BP1.

**Fig. 3:**
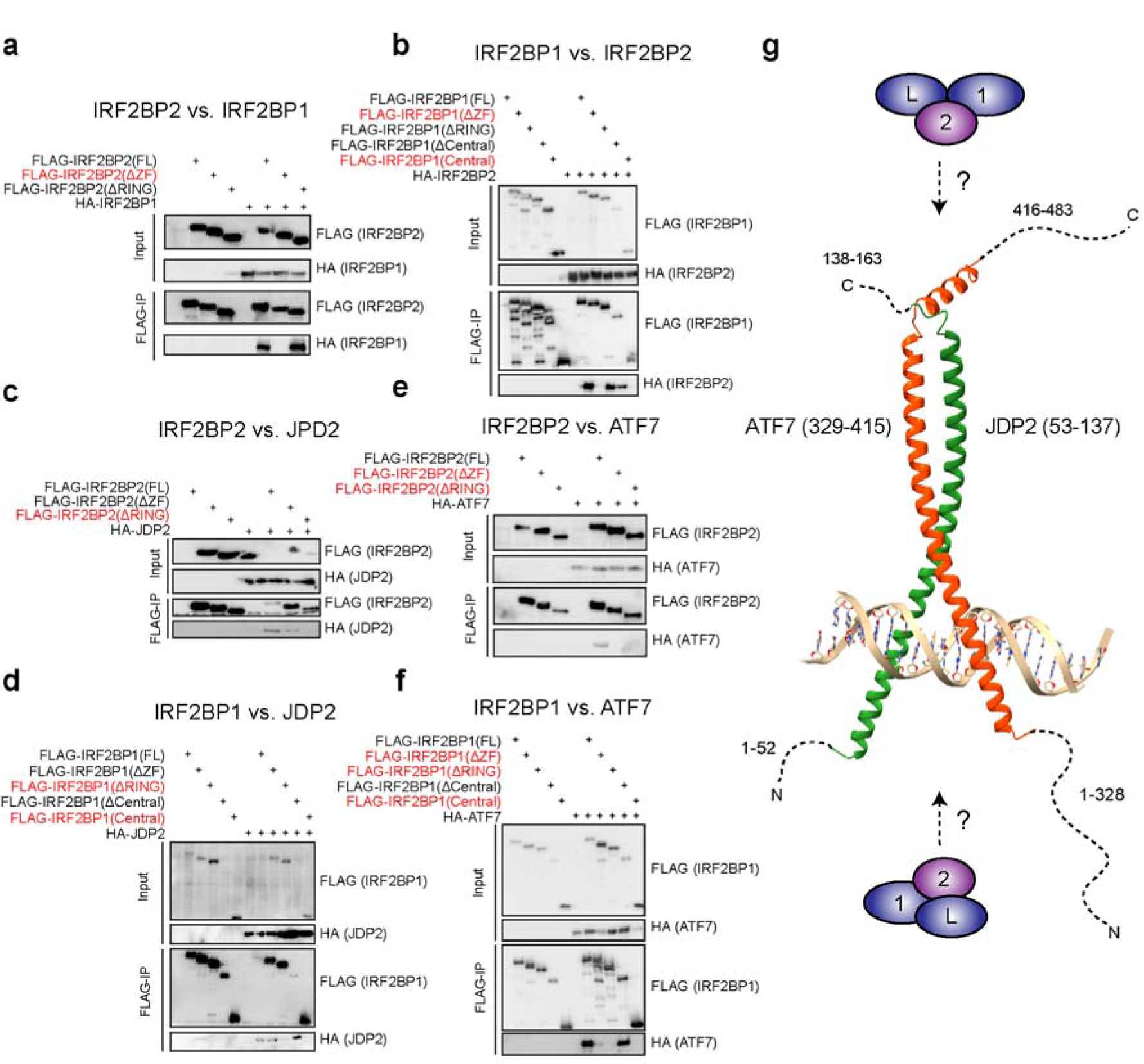
IRF2BP2 associates with ATF7 and JDP2 via multiple interaction sites. a) Western blot of a co-immunoprecipitation experiment of IRF2BP2 deletion mutants versus IRF2BP1. b) Western blot of a co-immunoprecipitation experiment of IRF2BP1 deletion mutants versus IRF2BP2. c) Western blot of a co-immunoprecipitation experiment of IRF2BP2 deletion mutants versus ATF7. d) Western blot of a co-immunoprecipitation experiment of IRF2BP2 deletion mutants versus JDP2. e) Western blot of a co-immunoprecipitation experiment of IRF2BP1 deletion mutants versus ATF7. f) Western blot of a co-immunoprecipitation experiment of IRF2BP1 deletion mutants versus JDP2. g) AlphaFold predicted dimerization of ATF7 and JDP2^25, 26^ modeled onto DNA using PDB: 1FOS^27^. All co-immunoprecipitation experiments were performed upon ectopic expression in HEK293 cells. Representative results of at least two biological replicates are shown. Red proteins indicate those that do not interact in the respective experiment.

For the interaction with JDP2, the C-terminal RING domains are required, for both IRF2BP1 and IRF2BP2, while the absence of the other globular domains still allows the interaction (**Fig. 3c, d**). Surprisingly, for optimal interaction with ATF7, the N-terminal zinc finger, and the C-terminal RING domains are necessary for both IRF2BP1 and IRF2BP2 (**Fig. 3 e, f**). This observation suggests that the interaction with ATF7 is facilitated via multiple interactions, which is supported by the fact that ATF7 is much larger than JDP2 (**Fig. 2e**). Notably, AlphaFold2 predicts the ATF7/JDP2 dimerization with high confidence (**Fig. 3g**), but it fails to identify convincing interaction sites for IRF2BP2 or IRF2BP1, suggesting a non-trivial interaction between these proteins. To assess whether perhaps the association between IRF2BP2 and IRF2BP1 is required for the interactions with ATF7 and JDP2, we obtained IRF2BP1 KO HEK293 cells (**Supplementary Fig. 4a**). Using these cells, we found that the absence of IRF2BP1 does not strongly impair the interactions of IRF2BP2 with ATF7 or JDP2, suggesting that the dimerization of IRF2BP2 and IRF2BP1 is not required (**Supplementary Fig. 4a**). Similar results were also obtained for the opposite experiment, where we tested the interaction of IRF2BP1 with ATF7 and JDP2, in the absence of IRF2BP2 (**Supplementary Fig. 4b**). However, this experiment does not rule out the possibility that the remaining IRF2BPL protein can compensate for the loss of IRF2BP1 or IRF2BP2. Another possible mechanism for interaction may involve SUMOylation of IRF2BP2, given that a SUMOylation-deficient mutant of IRF2BP2 (K566R)^13^ interacts less with ATF7, while the interaction with JDP2 is unaffected (**Supplementary Fig. 4c**).

Thus, although our experiments identified the crucial domains of IRF2BP2 and IRF2BP1 for the interaction with ATF7 and JDP2, more research will be required to clarify how exactly this association is established.

### ATF7 chromatin binding in THP1 cells depends on JDP2

Our biochemical experiments suggest that ATF7 and JDP2 form a heterodimer in AML cells that likely binds to DNA, similar to other AP-1 dimers^23^. The transcriptional activity of the ATF7/JDP2 dimer may be modulated by its association with the IRF2BP2 corepressor complex. To explore the relationship of these proteins in transcriptional regulation in more detail, we first investigated whether ATF7 relies on JDP2 for chromatin binding. We established ATF7 knockout (KO) and JDP2 KO THP1 cells, confirmed by Western blotting experiments (**Fig. 4a**). These cells did not exhibit any significant changes in proliferation (**Supplementary Fig. 3d**). In a ChIP-qPCR experiment, ATF7 binding was lost in the ATF7 KO cells, confirming the successful knockout of ATF7 (**Fig. 4b**). Interestingly, deletion of JDP2 also resulted in a substantial reduction of ATF7 binding (**Fig. 4b**), indicating that the absence of JDP2 impairs the chromatin binding of ATF7. Our attempts to also chromatin-immunoprecipitate JDP2 were not successful, preventing us from investigating the chromatin binding of JDP2 in further detail.

**Fig. 4:**
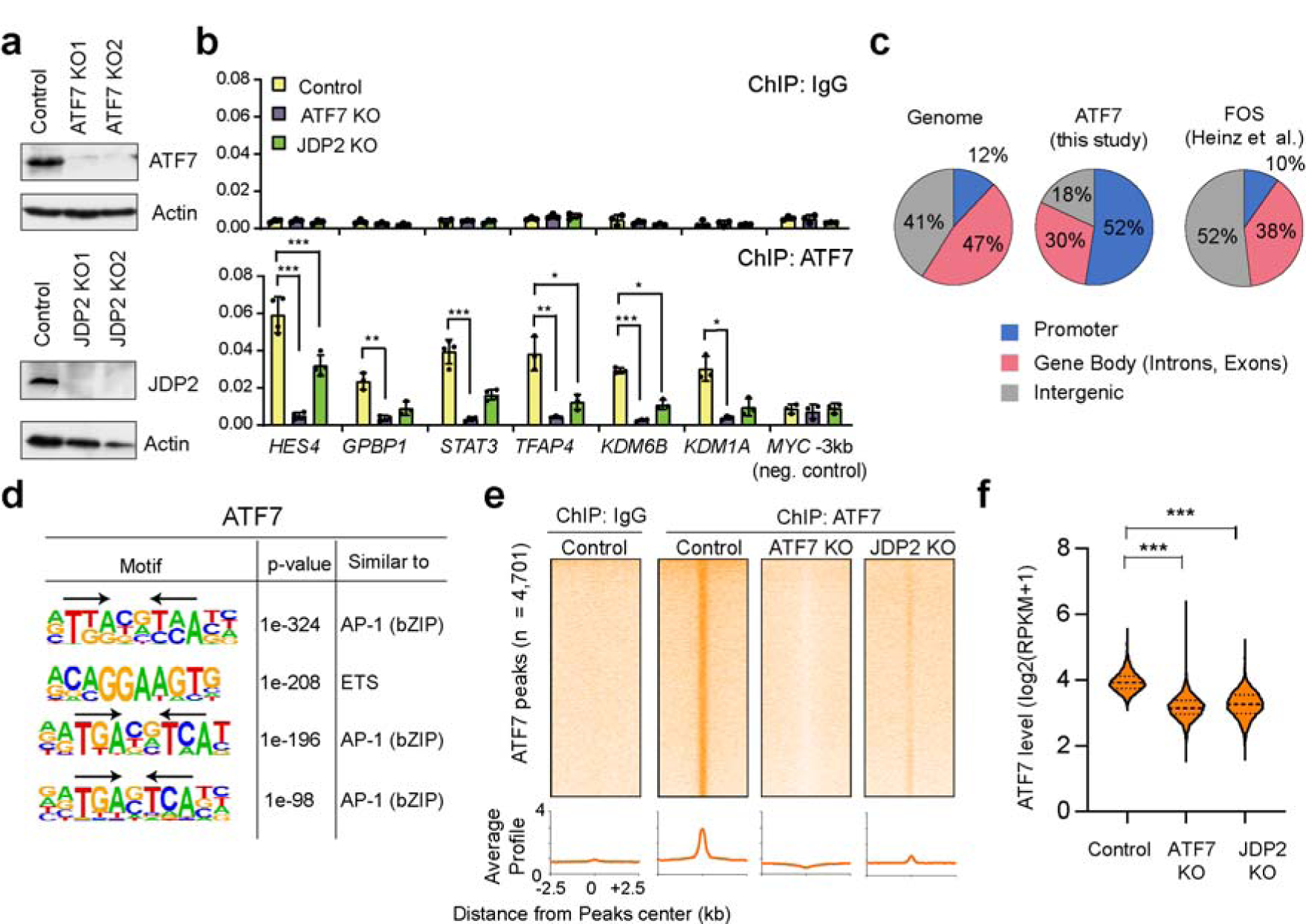
ATF7 chromatin binding requires JDP2. a) Western blot of THP1 cells with CRISPR/Cas9-mediated knockouts of ATF7 and JDP2. b) ChIP-qPCR of ATF7 in control, ATF7 KO, and JDP2 KO cells. Data represent the mean ± s.d. of at least three biological replicates. The significance was evaluated via a two-tailed unpaired Student’s t-test. c) Genomic distribution of ATF7 compared to FOS^28^. d) Motifs at ATF7 binding sites, identified via HOMER^30^. e) Heatmap of ATF7 ChIP-Seq results in control, ATF7 KO, and JDP2 KO cells. f) Violin plot showing the distribution of ATF7 ChIP-Seq signals at ATF7 peaks. Significance was evaluated via a two-sided Kolmogorov-Smirnov test.

To investigate the chromatin association of ATF7 and the impact of JDP2 deletion on a genome-wide level, we performed ATF7 chromatin immunoprecipitation followed by sequencing (ChIP-Seq) experiments in established cell lines. In total, we identified 4,701 significant ATF7 peaks in wild-type cells. Unlike most other AP-1 factors, such as FOS^28^, ATF7 showed predominant enrichment at promoters (**Fig. 4c**), suggesting a distinct transcriptional role for ATF7 compared to classical AP-1 transcription factors. Motif analysis revealed that the ATF7-bound locations were enriched for palindromic sequences resembling AP-1 binding sites (**Fig. 4d**). Notably, the most enriched motif consisted of a palindromic TTA sequence with a two-nucleotide spacing. This contrasts with the classical AP-1 binding motif, the TPA-responsive element (TRE), which consists of a palindromic TGA sequence with a one-nucleotide spacing^23^. This finding suggests a potential alternative DNA binding preference for ATF7/JDP2. Additionally, we observed a high enrichment of motifs for ETS transcription factors at the ATF7-bound locations, which is consistent with the known cooperation between AP-1 and ETS factors at target genes^29^.

In line with the qPCR experiments (**Fig. 4b**), ATF7 ChIP-Seq peaks were completely absent in the ATF7 KO cells and reduced to approximately 20% in the JDP2 KO cells (**Fig. 4e, f**). Interestingly, hardly any ATF7 peaks retained the ATF7 binding level in the JDP2 knockout cells (**Fig. 4e**), suggesting that JDP2 is the main partner of ATF7 in THP1 cells and is required for optimal ATF7 chromatin binding.

### Optimal IRF2BP2 chromatin binding requires the presence of ATF7 and JDP2

To investigate whether IRF2BP2 binds to similar locations as ATF7, we performed IRF2BP2 ChIP-Seq experiments in THP1 wild-type cells. We identified 21,518 significant peaks, which is more than previously published ChIP-Seq results of IRF2BP2 in MV4-11 and PDX1601 AML cells (**Fig. 5a, Supplementary Fig. 5a, b**), supporting the high quality of our data. Nonetheless, the previous ChIP-Seq data showed a high degree of overlap with our ChIP-Seq results (**Fig. 5a**), suggesting that the target genes of IRF2BP2 are similar in various AML cells. Similar to ATF7 (**Fig. 4c**), IRF2BP2 was also enriched at promoters (**Fig. 5b**).

**Fig. 5:**
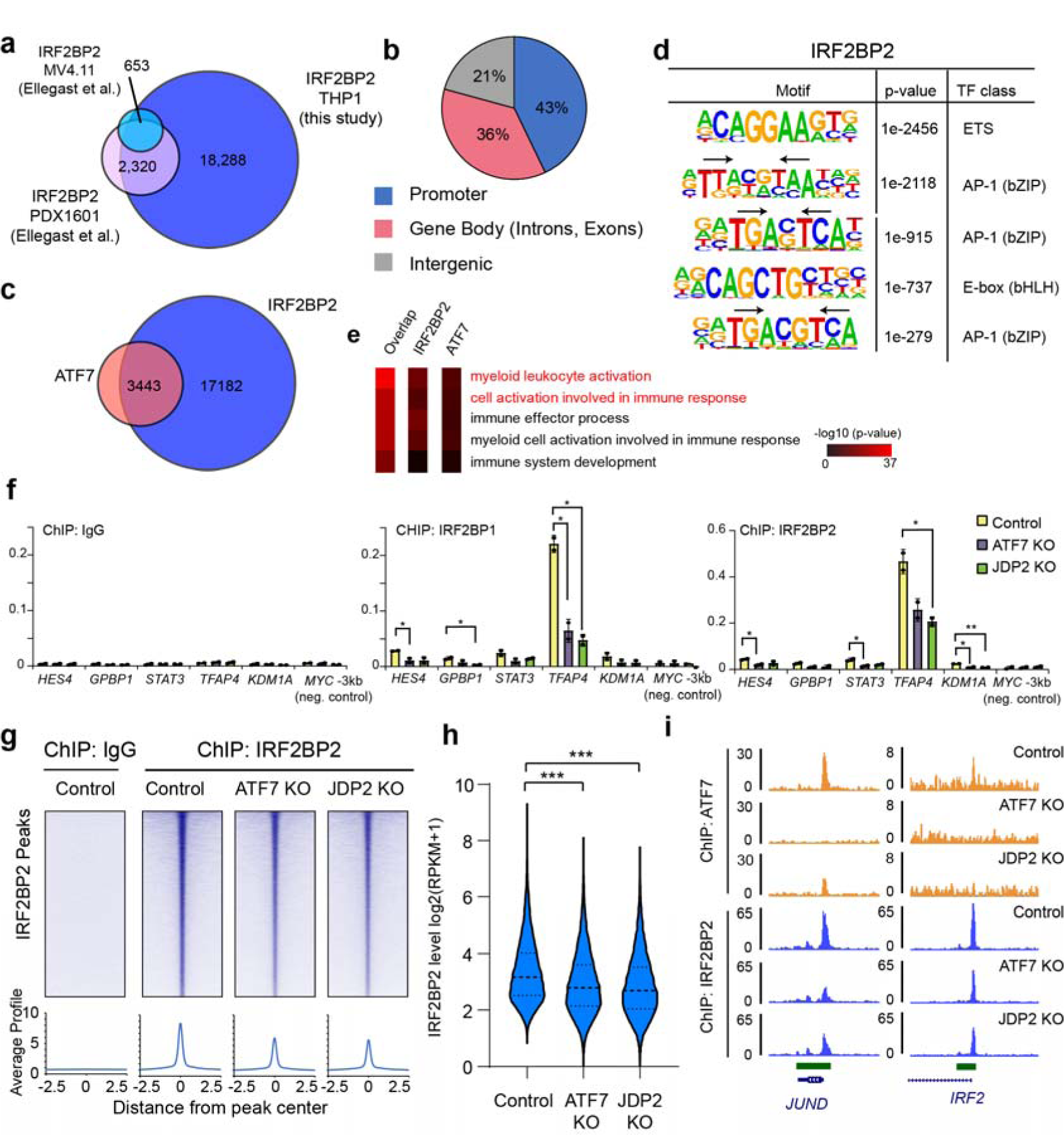
IRF2BP2 chromatin binding is reduced in the absence of ATF7 and JDP2. a) Venn diagramm showing overlap of IRF2BP2 ChIP-Seq peaks with IRF2BP2 ChIP-Seq data from Ellegast et al.^5^. See also Supplementary Fig. 4a and b. b) Genomic distribution of IRF2BP2. c) Venn diagram showing the overlap of IRF2BP2 peaks with ATF7 ChIP-Seq peaks. d) Motifs enriched at IRF2BP2 binding sites, identified via HOMER. e) Enriched gene ontologies at IRF2BP2/ATF7 overlapping peaks, and at ATF7 and IRF2BP2 peaks. Analysis performed via GREAT^31^. f) ChIP-qPCR experiments of IRF2BP1 and IRF2BP2 upon ATF7 and JDP2 knockout. Data represent the mean ± s.d. of at least two biological replicates. The significance was evaluated via a two-tailed unpaired Student’s t-test. g) Heatmaps showing IRF2BP2 ChIP-Seq signals at significant IRF2BP2 peaks upon ATF7 and JDP2 knockout. h) Violin plot showing IRF2BP2 levels at IRF2BP2 peaks upon ATF7 and JDP2 knockout. Significance was evaluated via a two-sided Kolmogorov-Smirnov test. i) Genome browser image showing the reduced levels of IRF2BP2 at the JUND and IRF2 genes upon ATF7 and JDP2 knockout.

Approximately 80% of the ATF7 peaks overlapped with the IRF2BP2 peaks (**Fig. 5c**), indicating a substantial binding overlap between the two proteins. Further analysis revealed that approximately 90% of the IRF2BP2 binding sites had some level of ATF7 binding (**Supplementary Fig. 5c**), demonstrating that IRF2BP2 commonly colocalizes with ATF7 in THP1 cells. Moreover, motif enrichment analysis revealed that the IRF2BP2 binding sites possess motifs similar to those of the ATF7 binding sites (**Fig. 4d, 5d**), further supporting their co-association at chromatin. Gene ontology analysis using the GREAT tool^31^ revealed that the target genes of ATF7 and IRF2BP2 are primarily associated with myeloid leukocyte activation and the immune response (**Fig. 5e**), consistent with the hypothesis that they play a crucial role in regulating genes involved in myeloid cell function.

Next, we asked to what extent the ATF7/JDP2 dimer is relevant for the chromatin binding of the IRF2BP2 complex. In ChIP-qPCR experiments, the deletion of ATF7 and JDP2 led to an approximately 50% reduction in IRF2BP2 at the investigated genes (**Fig. 5f**). Similar results were also obtained for IRF2BP1 (**Fig. 5f**), supporting that IRF2BP1 also requires ATF7/JDP2 for chromatin binding. Consistent with the ChIP-qPCR results, IRF2BP2 chromatin binding was globally reduced in both ATF7 and JDP2 KO cells (**Fig. 5g, h**). At about 5% of the loci, such as at the JunD gene (**Fig. 5i**), we observed a strong reduction in both knockouts (**Supplementary Fig. 5d, f**), suggesting that they are particularly dependent on the ATF7/JDP2 dimer. A total of 10 to 15% of the target genes were predominantly affected by either ATF7 or JDP2 KO, respectively (**Supplementary Fig. 5d, f**), raising the possibility that at these locations the absence of either ATF7 or JDP2 can be compensated by the formation of alternative AP-1 dimers. At most of the peaks (55%), such as at the IRF2 promoter (**Fig. 5i**), we observed an intermediate reduction in both knockouts (**Supplementary Fig. 5d, f**). Approximately 15% of IRF2BP2 peaks were unaffected (**Supplementary Fig. 5d, f**). These different groups show slightly different motif enrichments (**Supplementary Fig. 5e**), supporting the idea that they are targeted by a distinct set of transcription factors, which may explain the different consequences on IRF2BP2 chromatin binding upon ATF7 and JDP2 knockout. Notably, almost none of the IRF2BP2 peaks showed a reduction to the background level (**Fig. 5g, h**), implying that IRF2BP2 chromatin binding only partially depends on the ATF7/JDP2 dimer.

Our mass spectrometry analysis did not identify any additional strong candidate transcription factors that may contribute to the chromatin association of IRF2BP2 in AML cells, apart from the ATF7/JDP2 dimer (**Fig. 2b**). However, consistent with a previous study^9^, we observed a modest but non-significant enrichment of the NCOR1 and NCOR2 proteins (**Fig. 2b**). This finding supports the possibility that the IRF2BP2 may indirectly associate with chromatin through its interaction with the NCoR/SMRT corepressor complexes. In addition, we speculated that the formation of the IRF2BP2 complex, comprising IRF2BP2, IRF2BP1, and IRF2BPL, could be relevant for chromatin binding of IRF2BP2. To investigate this idea, we generated IRF2BP1 knockout THP1 cells. However, in ChIP-qPCR experiments, we observed no substantial reduction in the chromatin binding of IRF2BP2 in the absence of IRF2BP1 (**Supplementary Fig. 5g**), suggesting that IRF2BP2 does not require IRF2BP1 for chromatin binding.

Taken together, our results demonstrate that the ATF7/JDP2 dimer significantly contributes to the chromatin binding of the IRF2BP2 complex, as evidenced by the reduced occupancy of IRF2BP2 at target genes in the absence of ATF7 and JDP2. The presence of residual binding and the observed differences in dependency at specific loci suggest the involvement of additional recruitment mechanisms.

### The absence of IRF2BP2 leads to the overactivation of inflammatory pathways

Gene ontology analysis of the binding sites of IRF2BP2 and ATF7 revealed strong enrichment of genes associated with myeloid activation and inflammatory response (**Fig. 5e**), suggesting their involvement in regulating these cellular programs in AML cells. This hypothesis is supported by our analysis of TCGA data, which indicated that IRF2BP2 and ATF7/JDP2 might have opposing roles in regulating these pathways (**Fig. 2f-h**). To investigate the consequences of removing these proteins from THP1 cells, we conducted RNA-Seq experiments. We utilized THP1 cells with IRF2BP2 knockdown using two distinct shRNA constructs (**Fig. 1f**) and our established ATF7 and JDP2 knockout cells (**Fig. 4a**).

Examining the differentially regulated genes (**Fig. 6a**), we observed the most pronounced effect after IRF2BP2 knockdown, with 1639 dysregulated genes, indicating substantial changes in the gene expression pattern (**Fig. 6a**). This stronger effect on gene expression upon IRF2BP2 depletion is consistent with the more severe phenotype and higher score in the CRISPR screens (**Fig. 1a**). Importantly, analysis of the top 250 target genes of IRF2BP2 revealed that IRF2BP2 target genes predominantly became upregulated upon IRF2BP2 knockdown (**Fig. 6b**). This confirms the known repressive role of IRF2BP2 on a genome-wide level and is consistent with the findings of Ellegast et al. (**Fig. 6b**)^5^. In contrast, the top target genes of ATF7 were predominantly downregulated (**Fig. 6c**), albeit to a lesser extent, indicating a gene-activating role of ATF7 in THP1 cells. This result is in line with the presence of a transcriptional activation domain within ATF7 (**Fig. 2e**).

**Fig. 6:**
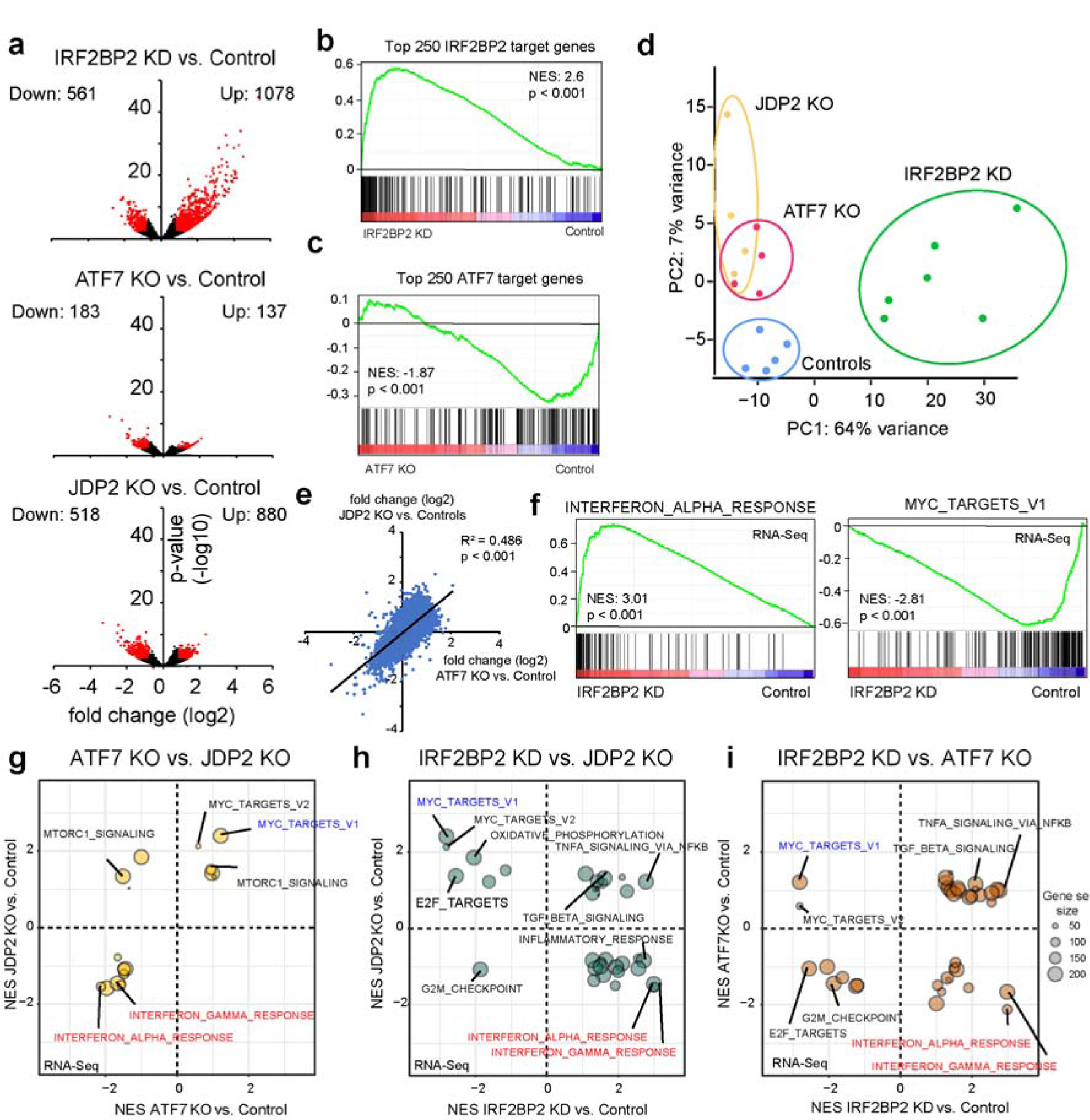
IRF2BP2 depletion activates inflammatory pathways in THP1 cells. a) Volcano plots showing significantly (fold change > 0.75, p-value < 0.01) differentially expressed genes upon knockdown of IRF2BP2 or knockout of ATF7 and JDP2. b) GSEA of the top 250 IRF2BP2 target genes and upon IRF2BP2 depletion. c) GSEA of the top 250 ATF7 target genes and upon ATF7 deletion. d) Principal component analysis of the RNA-Seq data. The data for IRF2BP2 were obtained from two independent shRNAs (3 replicates each). Two independent KO clones for ATF7 and JDP2 were analyzed (2 replicates each). e) Correlation of gene expression changes upon ATF7 and JDP2 KO. Significance was evaluated via ANOVA. f) GSEA plot of the top 2 hallmarks affected upon IRF2BP2 depletion in THP1 cells. Compare to Fig. 1i. g) Bubble plot comparing the GSEA results from ATF7 knockout and JDP2 knockout. h) Bubble plot comparing the GSEA results from IRF2BP2 knockdown and JDP2 knockout. i) Bubble plot comparing the GSEA results from IRF2BP2 knockdown and ATF7 knockout. NES = Normalized Enrichment Score, KO = Knockout, KD = Knockdown.

Consistent with these divergent effects on gene expression, principal component analysis (PCA) of the RNA-Seq data demonstrated distinct clustering of the IRF2BP2 knockdown cells (**Fig. 6d**). In contrast, ATF7 and JDP2 knockout cells exhibited close proximity (**Fig. 6d**), suggesting similar transcriptional effects. Indeed, further analysis revealed a strong correlation in the gene expression changes upon knockout of ATF7 and JDP2 (**Fig. 6e**), supporting our previous conclusion that these two proteins form a functional unit in these cells and that removing one of the two proteins also impairs the function of the other. For simplicity, we will refer to the knockouts of ATF7 and JDP2 as ATF7/JDP2 knockout in the following section.

To gain further insights into the specific pathways affected by IRF2BP2 knockdown, we conducted gene set enrichment analysis (GSEA). In the IRF2BP2 knockdown cells, the interferon alpha and gamma response pathways were the most significantly affected, exhibiting substantial upregulation (NES = 3.01 and 2.95, respectively) (**Fig. 6f**). Concurrently, MYC target genes were strongly downregulated (NES = −2.81) (**Fig. 6g**), consistent with the observed reduction in proliferation (**Fig. 1g**). Notably, the interferon pathway genes showed stronger occupancy by IRF2BP2 compared to MYC target genes (**Supplementary Fig. 6a**), supporting that interferon pathway genes are more directly affected. These two pathways were previously identified by us as significantly different between low and high IRF2BP2 expressing AML samples (**Fig. 1h-j**), thus demonstrating a high correlation between our RNA-Seq data and the TCGA data (**Supplementary Fig. 6b**). To validate the impact of IRF2BP2 knockdown on interferon response genes in other AML cell lines, we conducted RT-qPCR analysis on MV4-11 cells after IRF2BP2 knockdown. We found that similar to THP1 cells, most inflammatory genes including *IRF2*, *JUND*, and *IFIT2*, were strongly upregulated in MV4-11 cells (**Supplementary Fig. 6c**). These findings also align with a study by Ellegast et al., who investigated the consequences of IRF2BP2 depletion in MV4-11 AML cells using a dTAG system^5^. A direct comparison of the GSEA results of our RNA-Seq data from THP1 cells and the data from Ellegast et al. from MV4-11 cells showed a high degree of correlation (**Supplementary Fig. 6d**). Notably, Ellegast et al. proposed that the “TNFA_SIGNALING_VIA_NFKB” pathway is the most strongly affected pathway regulated by IRF2BP2. However, reanalysis of their data suggests that similar to our experiments, the interferon alpha and gamma response pathways were the most strongly altered hallmarks upon IRF2BP2 depletion (**Supplementary Fig. 6d**). This slight difference may stem from variances in bioinformatic pipelines. Nevertheless, the overall highly concordant results from our experiments in THP1 and MV4-11 cells, from Ellegast et al., and from the TCGA data strongly support the general conclusion that IRF2BP2 plays a critical role in controlling inflammatory pathways in AML cells.

Next, we investigated the consequences of ATF7 and JDP2 knockout. As expected, GSEA of the gene expression changes upon ATF7 and JDP2 knockout exhibited a high degree of similarity (**Fig. 6g**). Specifically, we found an upregulation of MYC target genes and the downregulation of interferon pathways upon their deletion (**Fig. 6g**). These findings are also largely consistent with the results obtained from the TCGA data (**Fig. 2f**) and are again opposite to the effects observed upon IRF2BP2 depletion. Consequently, the interferon pathways and MYC pathways anti-correlated between IRF2BP2 knockdown cells and ATF7/JDP2 knockout cells (**Fig. 6h, i**; lower right/upper left quadrant), similar to the TCGA data (**Fig. 2g, h**). Thus, the highly similar results obtained from the TCGA and our RNA-Seq data suggest that the anti-correlative property of the ATF7/JDP2-IRF2BP2 axis is a common theme in AML.

### IRF2BP2 knockdown enhances differentiation from myeloid progenitor cells

To gain insights into how changes in the expression level of IRF2BP2 may contribute to the development of AML during hematopoiesis, we investigated its role during differentiation from mouse myeloid progenitor cells, isolated from the bone marrow. IRF2BP2 and its interaction partners were detectable in the nucleus of cells isolated from the bone marrow (**Fig. 7a**). We knocked down IRF2BP2 in lineage-negative (Lin-) cells and performed colony formation assays in methylcellulose medium. The colony numbers obtained from the control cells and the knockdown cells were comparable (**Supplementary Fig. 7a**), but we found some morphological differences. We observed that the IRF2BP2 knockdown cells more often formed larger colonies (**Fig. 7b, c, Supplementary Fig. 7**) and that these colonies contained dendritic-shaped cells, which could not be detected in the control cells (**Fig. 7d**). Analysis of marker genes via RT-qPCR showed that several important differentiation markers were dysregulated in the IRF2BP2 knockdown cells (**Fig. 7e**). Specifically, we observed an increased expression of TGFβ, which plays a key role in differentiation processes^32^. Conversely, we observed reduced expression of the differentiation markers *Mmp9*, and *Ltf*. The stem cell marker *Kit* (c-kit) was also downregulated in the knockdown cells, supporting induction of differentiation (**Fig. 7e**). These changes in gene expression suggest an altered differentiation potential of the IRF2BP2 knockdown cells.

**Fig. 7.**
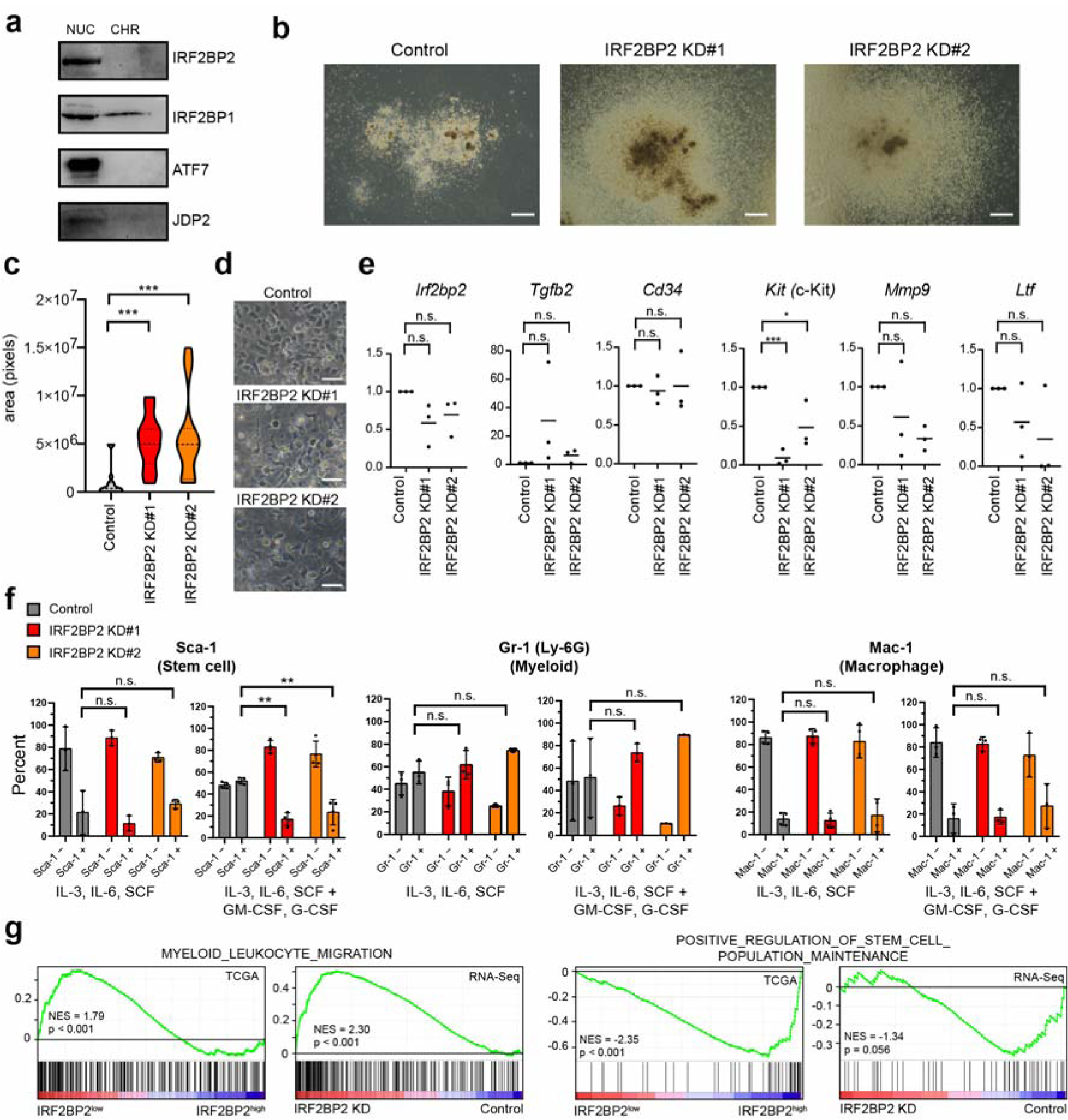
IRF2BP2 depletion influences differentiation from mouse bone marrow stem and progenitor cells. a) Western blot of components of the IRF2BP2 complex in cells isolated from mouse bone marrow. NUC = nucleoplasm, CHR = chromatin. b) Representative bright field photography of colonies (here CFU-GM) obtained upon colony formation assays of myeloid progenitor cells with and without IRF2BP2 KD in methylcellulose medium. Additional examples are shown in Supplementary Fig. 7b. Scale = 500 µm. c) Quantification of the area of colonies upon differentiation in methylcellulose. For each condition, 15 colonies were measured from two biological replicates. Significance was evaluated via a two-tailed unpaired Student’s t-test. d) Representative bright field microscopy photography of dendritic cell-shaped cells, seen in the IRF2BP2 knockdown cells compared to the control cells. Scale = 50 µm. e) RT-qPCR analysis of collected cells after colony formation assays. Data represents the mean of three biological replicates. Significance was evaluated via a two-tailed paired Student’s t-test. f) FACS quantification of surface makers after 10 days of differentiation of Lin-negative bone marrow cells in liquid culture. Data represent the mean +/− s.d. of at least three biological replicates. Significance was evaluated via a two-tailed unpaired Student’s t-test. Example histograms are shown in Supplementary Fig. 7b. g) GSEA of TCGA and our own RNA-Seq data regarding pathways involved in myeloid leukocyte migration and stem cell maintenance.

To better understand the role of IRF2BP2 during differentiation processes, we induced the wildtype and knockdown cells with IL-3, IL-6 and SCF and either in the presence or absence of G-CSF and GM-CSF, which stimulate myeloid differentiation^33^. Analysis of the cells after 10 days of liquid culture via flow cytometry suggests that in the absence of G-CSF and GM-CSF, the differentiation process was largely similar (**Fig. 7f**). In contrast, in the presence of G-CSF and GM-CSF, we observed a significantly decreased expression of the stem cell marker Sca-1 (Stem cell antigen-1) and an increased expression of Gr-1 (Ly-6G), a marker for myeloid differentiation, by trend (**Fig. 7f, Supplementary Fig. 7c**). No differentiation into B-cells or erythrocytes was observed under all conditions (**Supplementary Fig. 7d**). This observation suggests that the presence of IRF2BP2 inhibits the differentiation towards the myeloid lineage and favors a more stem cell-like state. The involvement of IRF2BP2 in differentiation processes is further supported by the expression data from TCGA and our RNA-Seq data. GSEA demonstrated that low expression of IRF2BP2 correlates with higher expression of genes involved in myeloid-related pathways, such as myeloid leukocyte migration and with lower expression of genes involved in stem cell maintenance (**Fig. 7g**). Consistent with the dendritic-shaped cells in the knockdown cells in the methylcellulose assays (**Fig. 7c**), we also observed an increased expression of dendritic cell differentiation pathway genes when IRF2BP2 is lowly expressed (**Supplementary Fig. 7e**).

Thus, we conclude that IRF2BP2 is likely a relevant regulator of hematopoietic stem cell differentiation processes. Its presence appears to favor stem cell-like characteristics and to inhibit differentiation towards the myeloid lineage and potentially monocytes-derived dendritic cells, which are also associated with inflammation. Consequently, IRF2BP2 upregulation in hematopoietic cells (**Fig. 1b**) may favor an undifferentiated state, which is typical for AML^1^.

## Discussion

Acute myeloid leukemia (AML) is a severe hematological malignancy characterized by the proliferation of undifferentiated myeloid progenitor cells^1^. Despite advances in treatment, AML remains a challenging disease to manage, and there is a need for more effective therapies. Inflammation plays a critical role in AML pathogenesis^4, 34^, and identifying the molecular mechanisms underlying this process is crucial for developing novel treatments. This study aimed to investigate the gene regulatory mechanisms of interferon regulatory factor 2 binding protein 2 (IRF2BP2) in AML, which has recently emerged as an important player in regulating inflammation in AML^5^, and which is required to maintain high proliferation of AML cells.

Using semi-quantitative mass-spectrometry, we identified the AP-1 heterodimer ATF7/JDP2 as a major interacting partner of IRF2BP2 (**Fig. 2b**). Previous work suggested that AP-1 transcription factors play a crucial role in setting up the transcriptional networks in AML^35^. ATF7 possesses a transactivation domain and typically functions as a transcriptional activator. In contrast, JDP2 is commonly described as a repressor^36^. To our knowledge, an AP-1 dimer established between ATF7 and JDP2 has not been described so far, suggesting that this combination is only established under certain circumstances. Furthermore, most other previously described transcription factors that have been described to interact with IRF2BP2 were not identified in our mass-spectrometry experiments. This observation raises the possibility that IRF2BP2 may interact with the ATF7/JDP2 dimer only in certain cell types, such as AML, while it may interact with other transcription factors in other contexts. If true, this ability may explain the rather versatile biological role of IRF2BP2 in development and diseases^37^. Our biochemical work implies that both the N-terminal zinc finger and the C-terminal RING domain are important for the optimal interaction of IRF2BP2 with ATF7/JDP2. We also revealed that the complex that consists of IRF2BP2, IRF2BP1, and IRF2BPL may not only hold together by the N-terminal zinc fingers but likely also involves the dimerizing central domains of IRF2BP1 and IRF2BPL. Currently, it remains unknown whether this domain may have additional functions and whether it is a subject of regulation. Future studies that clarify the function of this domain and the exact mechanism of how the IRF2BP2 complex interacts with the ATF7/JDP2 dimer may pave the way to develop strategies to perturb their gene regulatory functions.

On chromatin, ATF7 and IRF2BP2 strongly colocalize at many genes (**Fig. 5c**), and we demonstrated that the chromatin recruitment of IRF2BP2 is at least partially dependent on ATF7 and JDP2 (**Fig. 5 h, i**). This suggests that IRF2BP2 and ATF7/JDP2 form a functional unit on chromatin. Their target genes are linked to myeloid activation and inflammation (**Fig. 5e**). Consistently, depletion of either ATF7, JDP2, or IRF2BP2 significantly influences pathways related to inflammation, particularly interferon pathways (**Fig. 6f-h**). However, IRF2BP2 and ATF7/JDP2 affect gene transcription differently (**Fig. 6g-i**). While ATF7 and JDP2 act as positive regulators, IRF2BP2 functions as a negative regulator (**Fig. 8a-c**). The association of ATF7/JDP2 with the IRF2BP2 complex may therefore constitute an important regulatory hub to achieve a balanced level of intrinsic inflammation. A certain level of inflammation is critical for the optimal growth of AML cells^34^. On the other hand, too high levels of inflammation lead to cell death^5, 38^. Thus, the ATF7/JDP2-IRF2BP2 axis possibly represents a critical vulnerability in AML, and its dysregulation may contribute to AML development. By blocking the interaction of IRF2BP2 with ATF7/JDP2 or by disturbing the establishment of the IRF2BP2 complex, the optimal balance may be disrupted, leading to inflammatory overactivation, which ultimately leads to cell death (**Fig. 8b**). Potentially, interferon therapies that also induce inflammation in AML cells^38, 39^, may become more efficient when combined with approaches that target the IRF2BP2-ATF7/JDP2 axis.

**Fig. 8:**
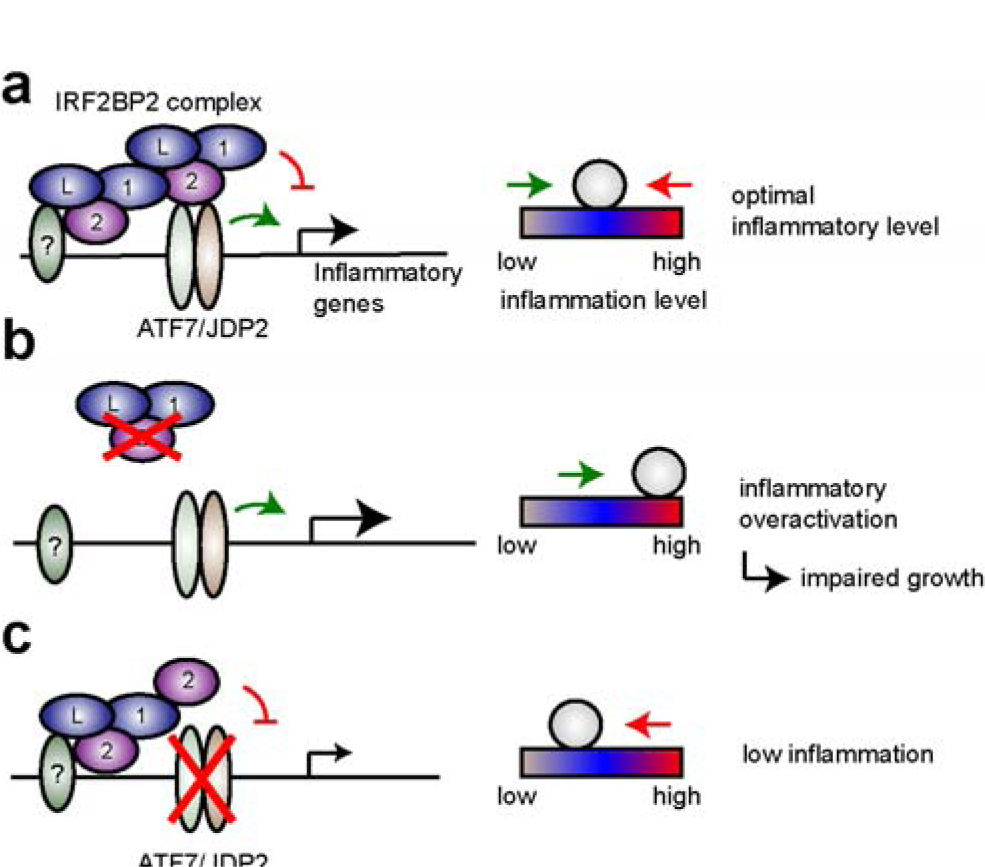
Model of the role of the ATF7/JDP2-IRF2BP2 axis in AML. a) ATF7/JDP2 activates inflammatory pathways, while IRF2BP2 is involved in counteracting ATF7/JDP2. This balanced regulation establishes an optimal level of inflammation required for high proliferation and rapid cancer progression. b) In the absence of IRF2BP2, inflammatory genes become strongly upregulated, leading to inflammatory overactivation and cell death. c) In the absence of ATF7/JDP2, their activating function is abolished. Due to alternative recruitment mechanisms, the repressive role of IRF2BP2 is partially retained, leading to a reduction in inflammatory pathways.

Inflammation has been proposed to play a key role in lineage decisions during hematopoiesis^40^. The delicate transcriptional balance established by the ATF7/JDP2-IRF2BP2 axis to regulate inflammation may therefore be an important aspect during this process. Our myeloid progenitor cell differentiation experiments support this idea (**Fig. 7**). Experiments in zebrafish also support the importance of IRF2BP2 for proper hematopoiesis^41^. More research will be required to understand the precise role of the ATF7/JDP2-IRF2BP2 gene regulatory system during hematopoiesis, and how its dysregulation may contribute to the development of AML.

In summary, our study provides insights into the molecular mechanisms of IRF2BP2 in AML and its interaction with ATF7/JDP2 to regulate inflammatory pathways. Our work suggests that the ATF7/JDP2-IRF2BP2 axis plays a crucial role in balancing inflammatory pathways in AML cells and possibly in other hematopoietic cells. This interplay, therefore, constitutes a potential vulnerability and offers a starting point to develop alternative strategies to treat AML patients.

## Methods

### Cell culture medium and handling

HEK293, HEK-T, THP1, and MV4-11 cells were grown at 37 °C with 5% (v/v) CO2. Growth media (RPMI for MV4-11 and THP1 cells, DMEM: F12/DMEM for HEK293 and HEK-T) were supplemented with FBS (10% v/v), penicillin (100 units/mL), and streptomycin (100 μg/mL).

### Virus production and knockdown of IRF2BP2

Cells were subjected to lentiviral infection to generate a knockdown/KO or OE of IRF2BP2 and its interactors in THP1 and MV4-11 cells. A virus was produced in HEK293-T cells by transfection of the packaging plasmids pMD2.g, psPAX2, and the respective pLKO.1 or LentiCrispr V2 vector, containing shRNA directed against control or IRF2BP2, respectively (shRNA sequences in **Supplementary Table 2**) with PEI solution (1 mg/mL). The virus-containing supernatant was collected 48 h after transfection. Cell lines were infected with the virus for 72 h. Then cells were selected with 1/0.5 μg/mL (THP1/MV4-11) of puromycin for 48 h. Subsequently, cells were counted and seeded at a density of 12 x 10^5^ in a six-well plate and collected after 48 h for RNAseq of KD. Puromycin selection was stopped at this time point. For proliferation experiments, cells were seeded at 1 x 10^4^ per well.

For the KO of ATF7, JDP2, and IRF2BP1/2 (guide sequences in **Supplementary Table 2**) in AML cell lines, cells were also lentivirally infected. After 48 h, the selection was started until a respective control mock infection was fully dead. Single clones were achieved by limiting dilution.

### Negative selection assay

The assay was grossly performed as described^42^. Briefly, THP1/MV4-11 cells were infected to stably express the CRISPR/Cas9 enzyme. The stable cell line was then transduced with sgRNA-expression vectors LRG (Lenti_sgRNA_EFS_GFP; Addgene plasmid #65656) with gRNAs for IRF2BP2, or a control gRNA, to achieve approximately 50% transduction rate. After 2 days of infection, the percentage of GFP sgRNA-expressing cells was measured using flow cytometry at different time points and normalized to the percentage of GFP+ cells on the starting day.

### Proliferation assay

Cell proliferation was measured with CellTiter-Glo® Luminescent Cell Viability Assay (Promega) according to the manufacturer’s instructions. Cells were measured every two days, up to seven days, with a plate reader Mithras LB 940 Multimode Plate Reader (Berthold Technologies). Values were normalized to day 0. The experiment was repeated three times.

### Antibodies

For the antibodies used, see **Supplementary Table 1**.

### Immunofluorescence staining

For immunofluorescence staining, HEK293 cells were seeded on coverslips. MV4-11 cells were attached overnight by coating the coverslips in poly-D-lysine (A003-E; Sigma Aldrich; 100µg/ml in H_2_O) for 1 h. THP1 cells were spun down onto the coverslips at 800g for 8 min. The cells were fixed with 4% formaldehyde (w/v), methanol-free (Thermo Fisher Scientific), and subsequently permeabilized with 0.5% Triton X-100 in PBS. Blocking was performed with 10% FBS in PBS. To detect ELOB and ELOC, the respective primary antibodies (**Supplementary Table 2**) were diluted at 1:500 in the blocking solution. After primary antibody incubation, the cells were washed three times with 0.5% Triton X-100 in PBS. Secondary antibody incubation was conducted using Alexa Fluor 488 and goat anti-rabbit IgG (H+L) (Thermo Fisher Scientific, Waltham, MA, USA; A-11008) at a 1:1,000 dilution. Following three washing steps, the coverslips were mounted onto microscopy slides using VECTASHIELD® Antifade Mounting Medium with DAPI (VECTOR Laboratories).

### Co-immunoprecipitation

HEK293 cells were seeded in 10-cm dishes at 2.2 × 10^7^ cells per dish. After 24 h, the expression constructs encoding for N-FLAG–tagged and/or N-Ha-tagged proteins were transfected using PEI. Details about the expression constructs are described in **Supplementary Table 5**. Two days after transfection, cell lysis was performed using Co-IP lysis buffer (50 mM Tris-Cl pH 7.5,150 mM NaCl, 1% Triton X-100, 1 mM EDTA, 10% glycerol, 1x protease inhibitor cocktail EDTA-free (Roche), and 0.5 mM PMSF). Cells were shaken for 30 min at 4 °C followed by centrifugation for 10 min at 13,000 rpm at 4 °C. The extract was used for FLAG-IP. Beads were equilibrated by washing three times with Co-IP lysis buffer. To bind FLAG-tagged proteins, extracts were incubated with 20 μL equilibrated anti-FLAG M2 Affinity Gel (Merck, A2220) for approximately 3 hours at 4 °C. After incubation, three washing steps with Co-IP wash buffer were performed (50 mM Tris-Cl pH 7.4, 250 mM NaCl, 1 mM EDTA, 1% Triton X-100, 10% glycerol). The FLAG beads were prepared for Western blotting through the addition of 2x Laemmli buffer. Detection of proteins in the input, supernatant, and IP fractions was conducted via Western blotting. Some co-immunoprecipitation experiments were performed semi-in-vitro, meaning that the proteins were first expressed separately in the cells, and united for the co-immunoprecipitation. Co-IP experiments were repeated at least two times.

### Mass-spectrometry

Nuclear extracts were prepared for a FLAG-IP to identify potential IRF2BP2 interaction partners via mass spectrometric analysis. For each co-immunoprecipitation experiment, 1-2 x 10^8^ THP1 cells stably expressing either FLAG-HA-IRF2BP2 (WT); or FLAG-HA-GFP as a control were used. After collection, cells were centrifuged at 2,000 rpm and 4 °C for 10 min. Cells were resuspended in 5-times packed cell volume (PV) hypotonic buffer (10 mM Tris-Cl pH 7.3, 10 mM KCl, 1.5 mM MgCl_2_ 0.2 mM PMSF, 10 mM ß-mercaptoethanol, 1x protease inhibitor cocktail tablet (Roche)) and shaken at 4 °C for 10-15 min. To remove cell debris, lysates were centrifuged at 3,500 rpm and 4 °C for 15 min. The pellet was resuspended in 1x pellet volume low salt buffer (20 mM Tris-Cl pH 7.3,20 mM KCl, 1.5 mM MgCl_2_, 0.2 mM EDTA, 25% glycerol, 0.2 mM PMSF, 10 mM ß-mercaptoethanol,1x protease inhibitor cocktail). Cells were dounced for 7-10x, then 0.66x pellet volume of high salt buffer (20 mM Tris-Cl pH 7.3, 1.2 M KCl, 1.5 mM MgCl_2_, 0.2 mM EDTA, 25% glycerol, 0.2 mM PMSF, 10 mM β-mercaptoethanol) was added gradually, while constantly mixing. The lysate was incubated while rocking for 30 min at 4 °C. It was then centrifuged at 13,300 rpm and 4 °C for 15 min and the supernatant was transferred to a “Slide-A-Lyzer” G2 Dialysis Cassette’ (3.5K) (Thermo Scientific) and dialyzed against 3 L of dialysis buffer (20 mM Tris-Cl pH 7.3, 100 mM KCl, 0.2 mM EDTA, 20% glycerol, 0.2 mM PMSF, 1 mM DTT) overnight. To perform the FLAG-IP, the material was retrieved from the dialysis chambers and centrifuged at 13,300 rpm and 4 °C for 30 min. The anti-FLAG M2 affinity gel (Sigma) was prepared by washing once with TAP buffer (50 mM Tris-Cl pH 7.9, 100 mM KCl, 5 mM MgCl_2_, 0.2 mM EDTA, 10% glycerol, 0.2 mM PMSF, 1 mM DTT), three times with 100 mM glycine (pH 2.5), once with 1 M Tris-Cl (pH 7.9) and finally once with TAP buffer. The extract was added to the beads and rotated at 4 °C for 4 h. Afterwards, the beads were washed three times with TAP buffer to wash away unbound material. From then on, two different protocols were employed depending on whether the material was to be analyzed by mass spectrometry or on a silver-stained gel. In the former case, the beads were washed three times in 50 mM ammonium bicarbonate (NH4HCO3). The IP-ed material was then sent in for mass-spectrometry analysis at the Biomedical Center Munich, protein analysis unit (Head: Axel Imhof), where it was analyzed using a Q Exactive HF-X Hybrid Quadrupole-Orbitrap Mass Spectrometer. If the material was to be loaded on a silver gel or used for Western blotting, the beads were incubated with 0.2 mg/mL single FLAG-peptide (Sigma) in TAP buffer and rotated at 4 °C for 1 h. Afterwards, the beads were centrifuged at 3,000 rpm at 4 °C for 5 min for elution. The supernatant, containing the immunoprecipitated material, was then used for downstream applications.

### Cellular Fractionation Assay

Fractionation assays were conducted with the “Subcellular Protein Fractionation Kit for Cultured Cells” (Thermo Fisher) and performed as advised by the manufacturer. The protein concentration was estimated, and equal amounts were loaded. FLAG as well as H3, Tubulin, KDM1A and SP-1, were detected via Western blotting (see **Supplementary Table 1** for antibodies).

### Myeloid progenitor cell isolation and infection

Male C57BL/6 mice aged approximately six to eight weeks were euthanized by cervical dislocation in accordance with the German animal welfare act (§ 4 (3) TierSchG). Front and hind leg bones were removed from the mice and cleared of remaining tissue. Bone marrow cells were collected by bisecting the bones and gentle extraction using short-pulse centrifugation and resuspended in PBS. In the first purification step, cells were separated on a gradient of PBS with lymphocyte separation media (LSM-A; Capricorn). To subsequently obtain murine stem cells and myeloid progenitor cells, cells were further selected using the Lineage Cell Depletion Kit (130-090-88; Miltenyi Biotec) using the autoMAC separator (Miltenyi Biotec) according to the manufacturer’s description. Collected lineage-negative (lin-) cells were incubated overnight in IMDM (12440-053; Gibco) with 20% FBS (10270-106; Gibco) and 1% Pen Strep (15140-122; Gibco). Additionally, the medium was enriched with the cytokines mIL-3 (10 ng/ml) (213-13; PeproTech), mIL-6 (50 ng/ml) (216-16; PeproTech) and mSCF (50 ng/ml) (250-03; PeproTech). On the following day, progenitor cells were lentivirally infected with shRNA constructs for mouse IRF2BP2 or a respective control, expressing GFP as a marker. After 48 h of infection, knockdown cells were sorted for GFP-positive signal. Afterwards, cells were seeded for differentiation or colony formation assays.

### Colony Formation Assay in Semi-Solid Medium

The cells were seeded in duplicates at 5x10^3^ in 1 ml of mouse methylcellulose complete media without Epo (HSC008; R&D Systems) containing mIL-3, mIL-6, and mSCF (concentrations as described above), on 35 mm dishes and cultivated at 37°C and 5% CO2. To prevent drying out, 35 mm dishes were placed in larger dishes filled with sterile water. Colonies were counted and photographed after 10 days. Methylcellulose medium was dissolved by adding IMDM medium and isolated cells were used for RNA preparation. Pictures were analyzed with ImageJ.

### Differentiation of myeloid progenitor cells

Cells were split and seeded in two different media. One half was incubated in IMDM with 20% FBS and 1% Pen Strep containing mIL-3, mIL-6 and mSCF (concentrations as described above) (“IL-3, IL-6, SCF”). The other half received additionally the stimulating factors mGM-CSF (20 ng/ml) (315-03; PeproTech) and mG-CSF (60 ng/ml) (250-05; PeproTech) (“IL-3, IL-6, SCF + GM-CSF, G-CSF”). Cells were cultivated at 37°C, 5% CO2 for 10 days and differentiation was subsequently evaluated with flow cytometric analyses and May-Grünwald-Giemsa staining.

### FACS

Differentiated cells were collected and washed twice with PBS. Subsequently, cells were subjected to a live/dead staining with the Zombie NIR™ Fixable Viability Kit (423105; BioLegend) and incubated in the dark for 10 min. Then, the cells were washed with MACS buffer (2 mM EDTA in PBS, 0,2% BSA) and co-stainings with APC-coupled and PerCP/Cy5.5-coupled antibodies was performed by incubation for 45 min at 4 °C in the dark. Stainings of surface proteins Sca-1, CD117 (c-kit), CD11b (Mac-1), Gr-1, CD19, and Ter-119 were conducted, including respective antigen isotype controls for the applied antibodies (for a detailed list of antibodies see **Supplementary Table S1**). Cells were washed twice with PBS before resuspension in MACS buffer for subsequent flow cytometric analysis using the CytoFLEX (Beckman Coulter).

### RT-qPCR and RNA-Seq

RNA isolation was performed using the “RNeasy Mini Kit” (74104; Qiagen) according to the manufacturer’s instructions. An additional on-column DNA digest (79254; Qiagen) was added between the washing steps. According to the manufacturer’s manual, the Tetro cDNA Synthesis Kit (BIO-65043; BioLine) was used for transcribing mRNA into cDNA. Subsequently, cDNA was diluted 1:20 with water for use in RT-qPCR. For analysis by real-time quantitative PCR, the MyTaq™ Mix was used. For gene expression analysis, values were normalized to GAPDH or Actin expression (primers in **Supplementary Table 3**).

RNA-Seq was performed by Biomarker Technologies (BMKGENE). Libraries were made using a Hieff NGS® Ultima Dual-mode mRNA Library Prep Kit for Illumina® and sequenced using an Illumina NovaSeq 6000.

### ChIP-qPCR and ChIP-Seq

Cells were seeded on 15 cm plates at 3 × 106 cells per plate to prepare chromatin and cultivated until reaching 70–90% confluence. First, 1% formaldehyde was added to the medium, and the plates were slowly swayed for 10 min to fix the cells. The fixation was stopped by adding 125 mM glycine for 5 min. Subsequently, the cells were washed twice with PBS and scraped in 1 mL of cold buffer B (10 mM HEPES/KOH, pH = 6.5; 10 mM EDTA; 0.5 mM EGTA; 0.25% Triton X-100) per 15 cm plate. All plates containing the same cell line were pooled in a 15 mL tube. The tubes were centrifuged for 5 min at 2,000 rpm and 4 °C. The supernatant was removed, and the pellet was resuspended in 1 mL of cold buffer C (10 mM HEPES/KOH, pH = 6.5, 10 mM EDTA, 0.5 mM EGTA, 200 mM NaCl) per 15 cm plate followed by a 15 min incubation time on ice. Then, the tubes were centrifuged with the same settings as mentioned before. After removing the supernatant, the pellet was resuspended in 200 µL of cold buffer D (50 mM Tris/HCl, pH = 8.0, 10 mM EDTA, 1% SDS 1xPIC (cOmplete™, Protease Inhibitor Cocktail; Roche; 04693116001)) per 15 cm plate, vortexed, and incubated for 10–20 min on ice. To shear the chromatin, the samples were sonicated two times for 7 min each using a precooled Bioruptor^®^ (Diagenode). The samples were centrifuged for 10 min at 13,000 rpm and 4 °C. The supernatant contained the sheared chromatin.

Chromatin immunoprecipitation (ChIP) for ChIP-qPCR was performed according to the one-day ChIP kit protocol (Diagenode; C01010080). For each ChIP, 3 µg of antibodies were used (see **Supplementary Table 1** for antibody information). The primers for ChIP-qPCR are presented in **Supplementary Table 4**.

To prepare the samples for ChIP-sequencing, the one-day ChIP kit protocol was used as described above, but the DNA purification was modified. For DNA elution, the beads were incubated with 230 µL elution buffer (100 mM NaHCO3, 1% SDS) for 30 min at room temperature while shaking. Afterwards, the samples were centrifuged at 13,000 rpm for 1 min, and 200 µL of supernatant was transferred to a fresh tube. The input DNA was dissolved in 50 µL of dH2O, and 150 µL of elution buffer was added to obtain an equal volume in all samples. Eight microliters of 5 M NaCl was added to each sample, and the samples were incubated at 65 °C overnight to reverse the cross-linking.

On the next day, 8 µL of 1 M Tris/Cl pH 6.5, 4 µL 0.5 M EDTA, and 2 µL of Proteinase K (10 µg/µL) were added to each sample. All samples were incubated at 45 °C for 1 h while shaking. The DNA was purified using the QIAquick PCR Purification Kit (Qiagen), whereby all samples prepared with the same antibody were pooled on the same column. To elute the DNA, the columns were incubated for 1 min with 30 µL of sterile 2 mM Tris/Cl, pH 8.5, and centrifuged at 13,000 rpm for 1 min.

The concentration of the samples was determined using the Quant-iT™ dsDNA Assay Kit (Thermo Fisher Scientific, Waltham, MA, USA; Q33120) and the NanoDrop™ 3300 (Thermo Fisher Scientific). At least 4 ng of DNA was used for library preparation. ChIP-Seq libraries were made using Microplex library preparation kit v2 (Diagenode) and sequenced using Illumina NextSeq 550 at the Genomics Core Facility Marburg (Center for Tumor Biology and Immunology, Hans-Meerwein-Str. 3, 35043 Marburg, Germany).

### Bioinformatics analysis

ChIP-Seq data were mapped to the human genome hg38 using bowtie^43^, allowing 1 mismatch. BigWig files were obtained using DeepTools/bamCoverage^44^. Significant peaks were obtained using Galaxy/MACS2^45^. The genomic distribution of the peaks was analyzed using ChIPSeeker. Heatmaps and profiles were created using Galaxy/DeepTools^44^. Motif analysis was performed using HOMER^30^. Gene ontology of ChIP-Seq data was performed using GREAT^31^. The top target genes were identified based on the ChIP-Seq signal at promoters.

RNA-Seq data were aligned to the human transcriptome (GenCode 43) using Galaxy/RNA-Star^46^. Differentially expressed genes were obtained using DeSeq2^47^. Genes with a log2-fold change of more than 0.75 and a p-value lower than 0.01 were considered significantly dysregulated. Gene set enrichment analysis was performed using GSEA software with standard settings^48^. Bubble plots were created using the results from GSEA (cut-off FDR p-value < 0.1) via ggplot2^49^.

The following internet databases and tools were used: Galaxy Europa^50^, DeepTools^44^, DepMap^51^, GREAT^31^, Bioconductor/R^52^, GSEA^48^, GePIA^21^, AlphaFold Database^24^, Google Colabs/AlphaFold2^26^, GDC Data Portal^19^ and Kaplan-Meier-Plotter^22^.

The following public data were used: ChIP-Seq of FOS in THP1 cells (GSM3112113)^28^, ChIP-Seq of IRF2BP2 in MV4-11 cells (GSM5155146, GSM5155147, GSM5155148)^5^, ChIP-Seq of IRF2BP2 in PDX1601 cells (GSM5908693)^5^. RNA-Seq upon IRF2BP2 degradation in MV4-11 cells (GSM5155162 to GSM5155167)^5^. RNA-Seq data from TCGA^19^.

### Statistical Analysis

Statistical analysis was performed as described in the figure legends. Error bars indicate standard deviation (s.d.). The significance of theqPCR results was analyzed via ANOVA or Student’s t-tests. The significance of the GSEA was evaluated by the GSEA software. The significance of the ChIP-Seq levels at IRF2BP2/ATF7 peaks was evaluated using a two-sided Kolmogorov-Smirnov test. All biological experiments were performed in at least in two replicates. RNA-Seq was performed with at least two replicates per condition.

### Data availability

All data relevant to the study are included in the paper or uploaded as supplementary information. Source data are provided with this paper. All data and sources associated with this study are available from the corresponding author upon reasonable request. RNA-Seq and ChIP-Seq data were deposited in the Gene Expression Omnibus (GEO) database. Mass-spectrometry data were uploaded to PRIDE.

## Acknowledgments

This project was supported by the German José Carreras Leukemia Foundation (DJCLS 06 R/2022), the Fritz Thyssen Foundation (10.20.1.005MN) and the German Research Foundation (DFG, 384027541, 416910386) to R.L. Flow cytometric experiments in this work were supported by a grant from the Deutsche Forschungsgemeinschaft (project 443291381).

We acknowledge the Protein Analytics Unit at the Biomedical Center, Ludwig-Maximilians University Munich, for providing services and assistance with data analysis. We thank Frauke Melchior (Heidelberg University) for providing IRF2BP2 plasmids. We thank the cellular imaging core facility at the University of Marburg, especially Katrin Roth for her advice. We thank the FACS facility at the University of Marburg, especially Hartmann Raifer and Heinke Lucks for their support and advice.

## Authors Contribution

S.F. performed most biochemical and biological experiments. L.M.W., B.S., M.F., C.S. and J.K. contributed experiments and/or material. A.N. performed next-generation sequencing. I.F. performed mass-spectrometry analysis. R.L. performed bioinformatics analyses. U.M.B, T.S., A.N and R.L. supervised the work. R.L. and S.F. conceived the study. R.L. and S.F. wrote the manuscript with input from all authors.

---

**Supplementary Fig. 1:**
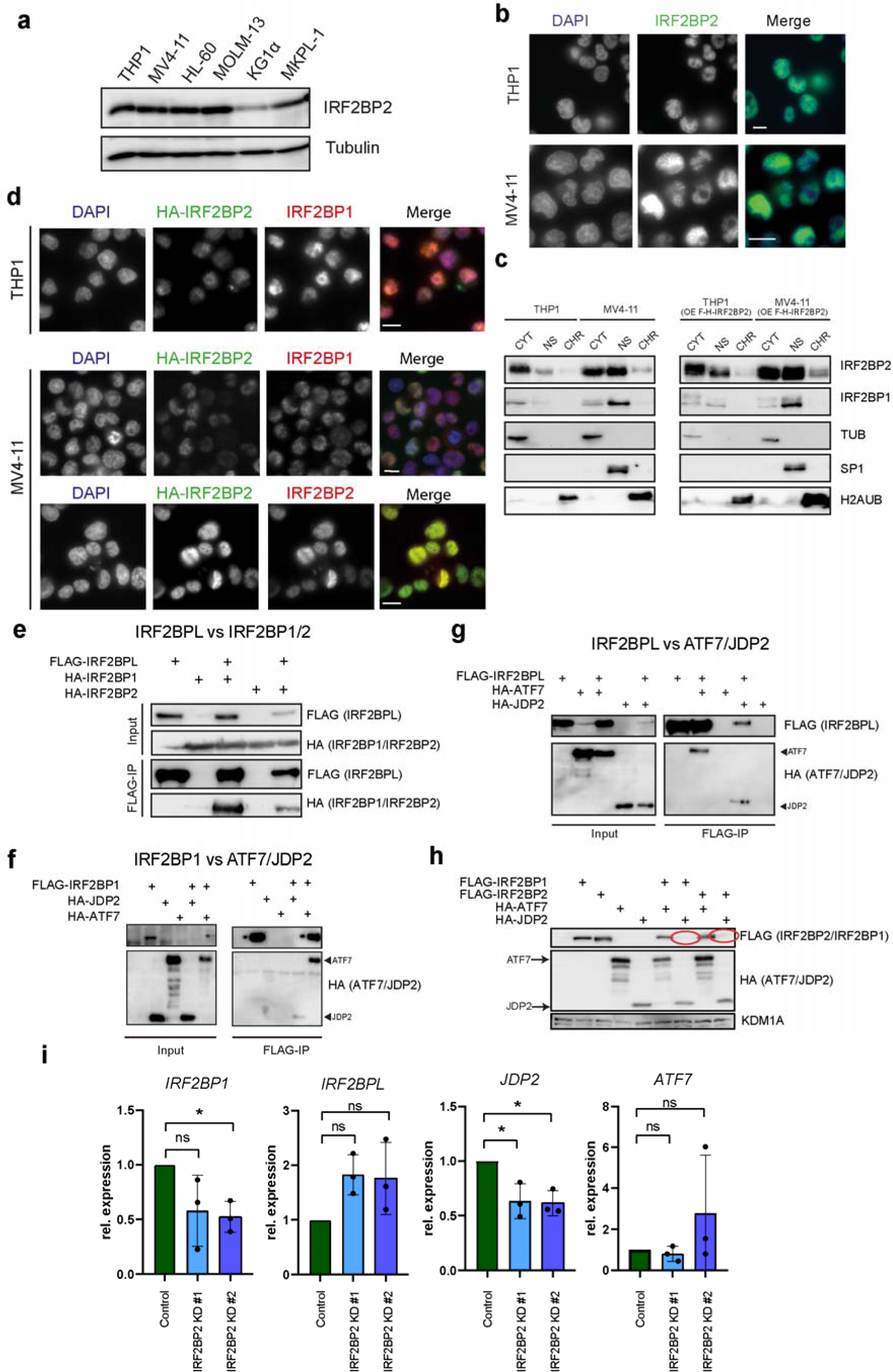
All members of the IRF2BP2 complex interact with ATF7 and JDP2. a) Western blot of endogenous IRF2BP2 in human AML cell lines. b) Immunofluorescence of endogenous IRF2BP2 in THP1 and MV4-11 cells. Scale bar = 10 µM. c) Immunofluorescence of HA-tagged IRF2BP2 in THP1 and MV4-11 cells. Costaining with IRF2BP1 shows colocalization in both cell lines. Scale bar = 10 µM. d) Cellular fractionation showing enrichment of IRF2BP2 in the nuclear and chromatin fraction of THP1 and MV4-11 cells. CYT = cytoplasm, NS = nucleoplasm, CHR = chromatin e) Co-immunoprecipitation experiments showing the interaction of IRF2BPL with IRF2BP1 and IRF2BP2. f) Co-immunoprecipitation experiments showing the interaction of IRF2BP1 with ATF7 and JDP2. g) Co-immunoprecipitation experiments showing the interaction of IRF2BPL with ATF7 and JDP2. h) The negative effect of JDP2 overexpression on the protein levels of IRF2BP1 and IRF2BP2. Red circles indicate positions where IRF2BP1 and IRF2BP2 are lacking upon JDP2 overexpression. i) RT-qPCR analysis of IRF2BP2 interaction partners upon IRF2BP2 knockdown in THP1 cells. Data represent the mean +/− s.d. of three biological replicates. Significance was evaluated via ANOVA.

**Supplementary Fig. 2.**
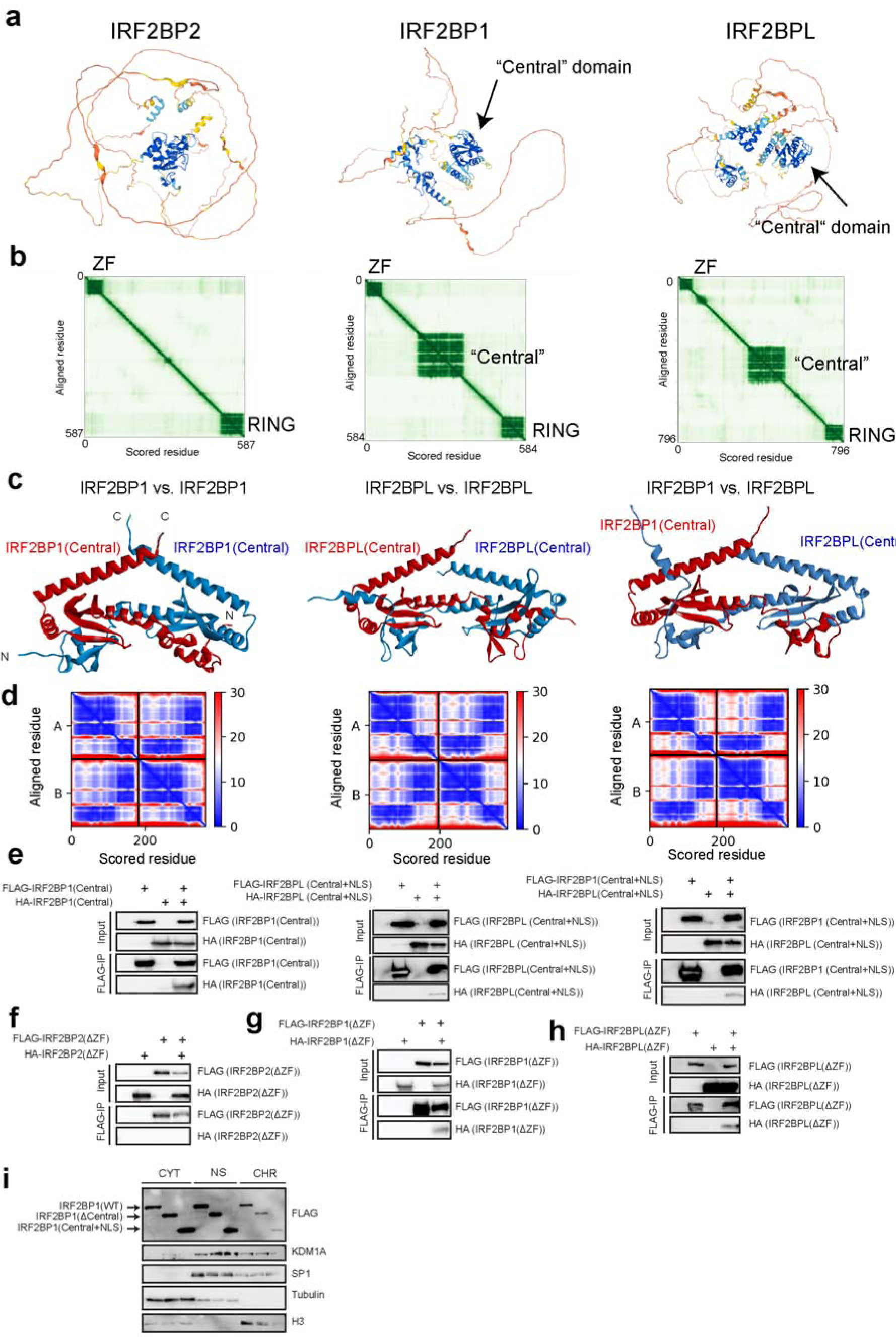
IRF2BP1 and IRF2BPL possess a central dimerization domain. a) AlphaFold2 predicted structures of IRF2BP2, IRFB2P1 and IRF2BPL obtained from the AlphaFold database^24^, of the human proteins (IDs: Q7Z5L9, Q8IU81, Q9H1B7) b) Alignment scores of the predictions from a). c) AlphaFold2-predicted^26^ association of two central domains, either of IRF2BP1, IRF2BPL or their combination. d) Alignment scores of the predictions from c). e) Co-immunoprecipitation experiments with the central domains, showing homo- and heterodimerization in vivo. f) Co-immunoprecipitation experiment of IRF2BP2 with itself without the zinc finger domain. g) Co-immunoprecipitation experiment of IRF2BP1 with itself without the zinc finger domain. h) Co-immunoprecipitation experiment of IRF2BPL with itself without the zinc finger domain. i) Cellular fractionation after ectopic expression of IRF2BP1 constructs in HEK293 cells.

**Supplementary Fig. 3:**
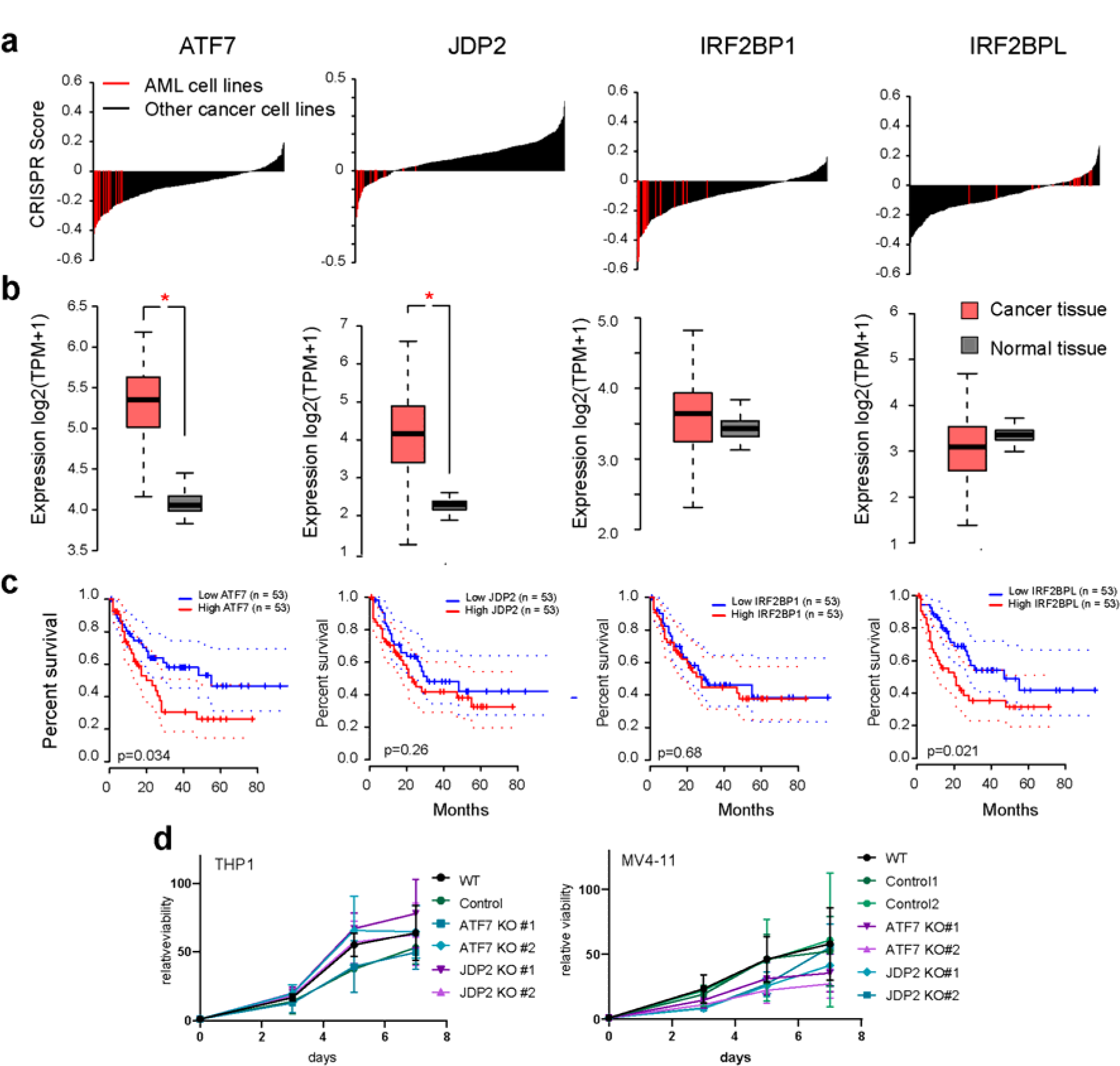
Analysis of IRF2BP2 interaction partners in AML. a) CRISPR scores of ATF7, JDP2, IRF2BP1 and IRF2BPL in AML (red) and non-AML cell lines, based on data from Meyer et al.^17^. b) Gene expression of the four genes in AML versus normal tissue, based on data from GePIA^21^. Red stars indicate significantly different expressions. c) Kaplan-Meier survival curve of AML patients based on ATF7, JDP2, IRF2BP1 and IRF2BPL expression. Data were derived from TCGA and were visualized using GePIA^21^. d) Proliferation of THP1 and MV4-11 AML cells upon knockout of ATF7 and JDP2. Data represent the mean +/− s.d of three biological replicates.

**Supplementary Fig. 4:**
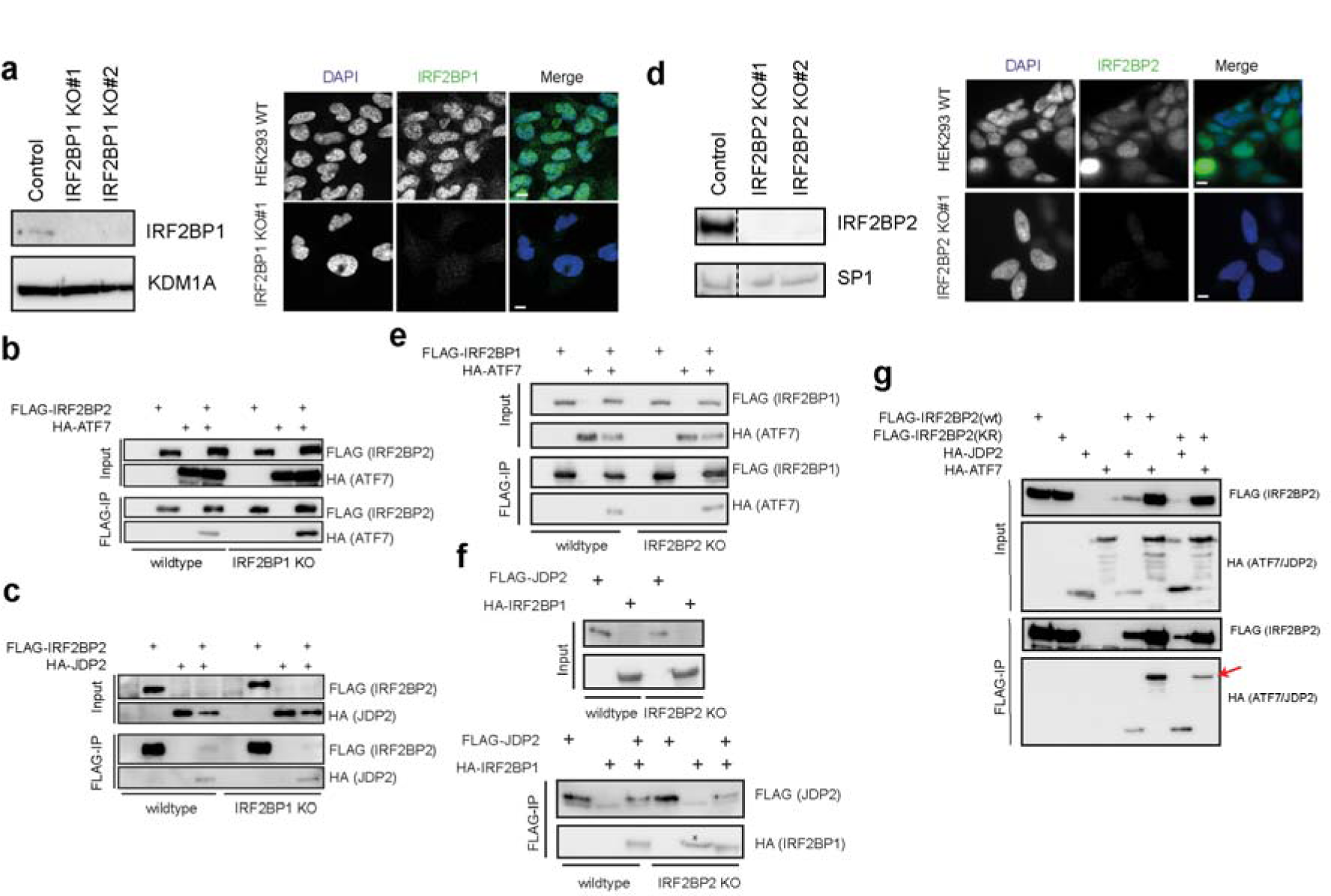
Co-immunoprecipitation experiments in IRF2BP1 or IRF2BP2 knockout cells. a) Western blot and immunofluorescence upon CRISPR-mediated knockout of IRF2BP1. Clone 1 was used for experiments. b) Western blot analysis of the co-immunoprecipitation of ATF7 with IRF2BP2 in IRF2BP1 KO cells. c) Western blot analysis of the co-immunoprecipitation of JDP2 with IRF2BP2 in IRF2BP1 KO cells. d) Western blot and immunofluorescence upon CRISPR-mediated knockout of IRF2BP2.Clone 1 was used for experiments. e) Western blot analysis of the co-immunoprecipitation of ATF7 with IRF2BP1 in IRF2BP2 KO cells. f) Western blot analysis of the semi-in vitro co-immunoprecipitation of JDP2 with IRF2BP2 in IRF2BP1 KO cells. Asterisks indicate an unspecific band. g) Effect of ubiquitination deficient IRF2BP2 on the interaction with ATF7.

**Supplementary Fig. 5:**
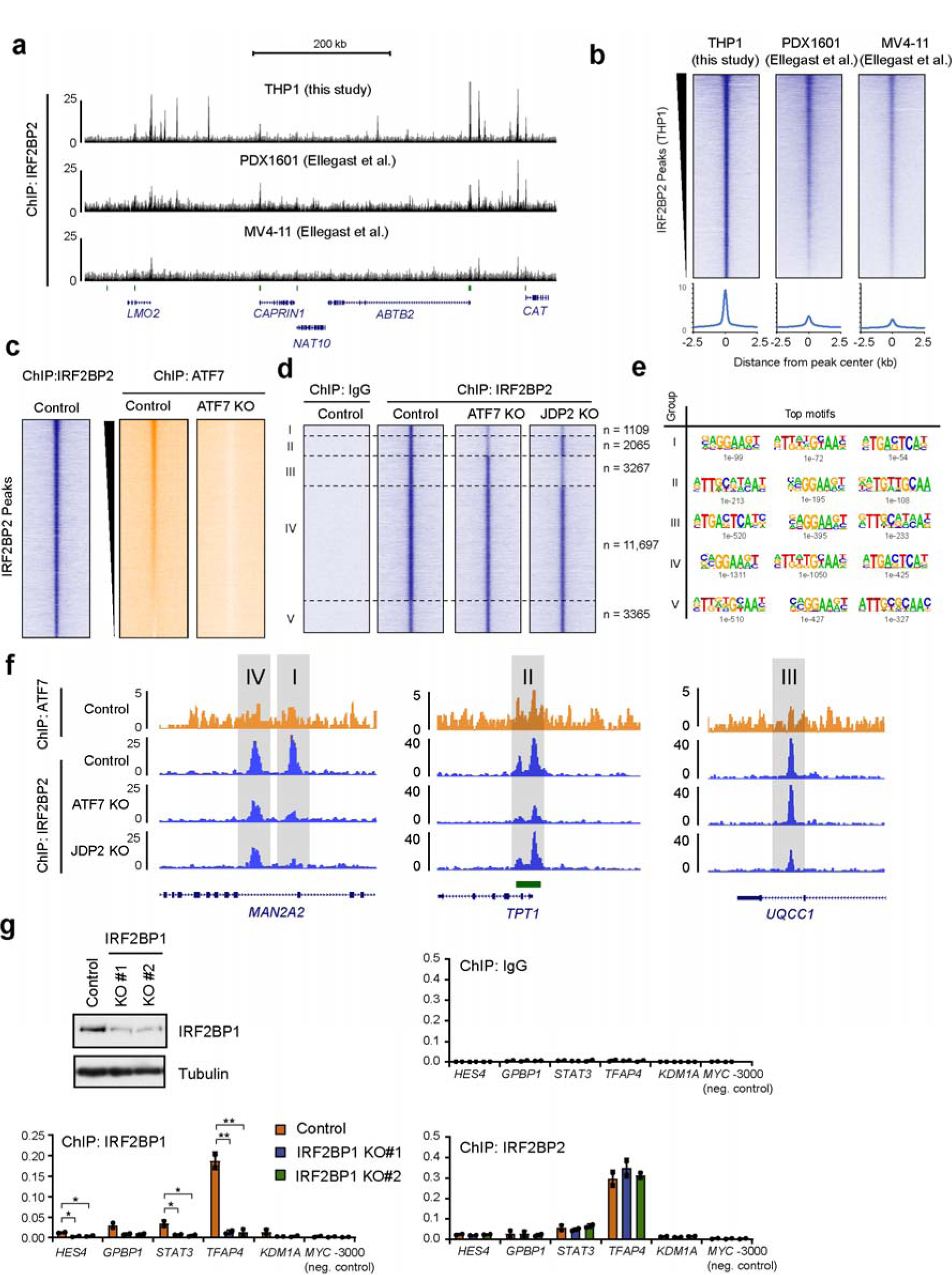
Additional analysis of IRF2BP2 ChIP-Seq results. a) Genome browser view showing the comparison of the ChIP-Seq results from this study, with those from Ellegast et al.^5^. b) Heatmap of the ChIP-Seq data from a). c) Heatmaps showing ATF7 ChIP-Seq signals in wildtype and ATF7 knockout cells at all IRF2BP2 peaks. Most IRF2BP2 peaks do possess some level of ATF7. d) Groups of IRF2BP2 peaks, categorized based on their reduction upon ATF7 and JDP2 knockout. Group I: Strongly reduced upon ATF7 and JDP2 knockout. Group II: Reduced mainly upon ATF7 knockout, Group III: Reduced mainly upon JDP2 knockout. Group IV: Slightly reduced in both knockouts. Group V: Not affected upon ATF7 or JDP2 knockout. e) Motifs enriched at the peaks from the five groups from d). f) Genome browser view of examples for groups I to IV from d). g) ChIP-qPCR in IRF2BP1 knockout THP1 cells, compared to control cells, using antibodies against IRF2BP1 and IRF2BP2. Data represent the mean ± s.d. of two biological replicates. The significance was evaluated via a two-tailed unpaired Student’s t-test.

**Supplementary Fig. 6:**
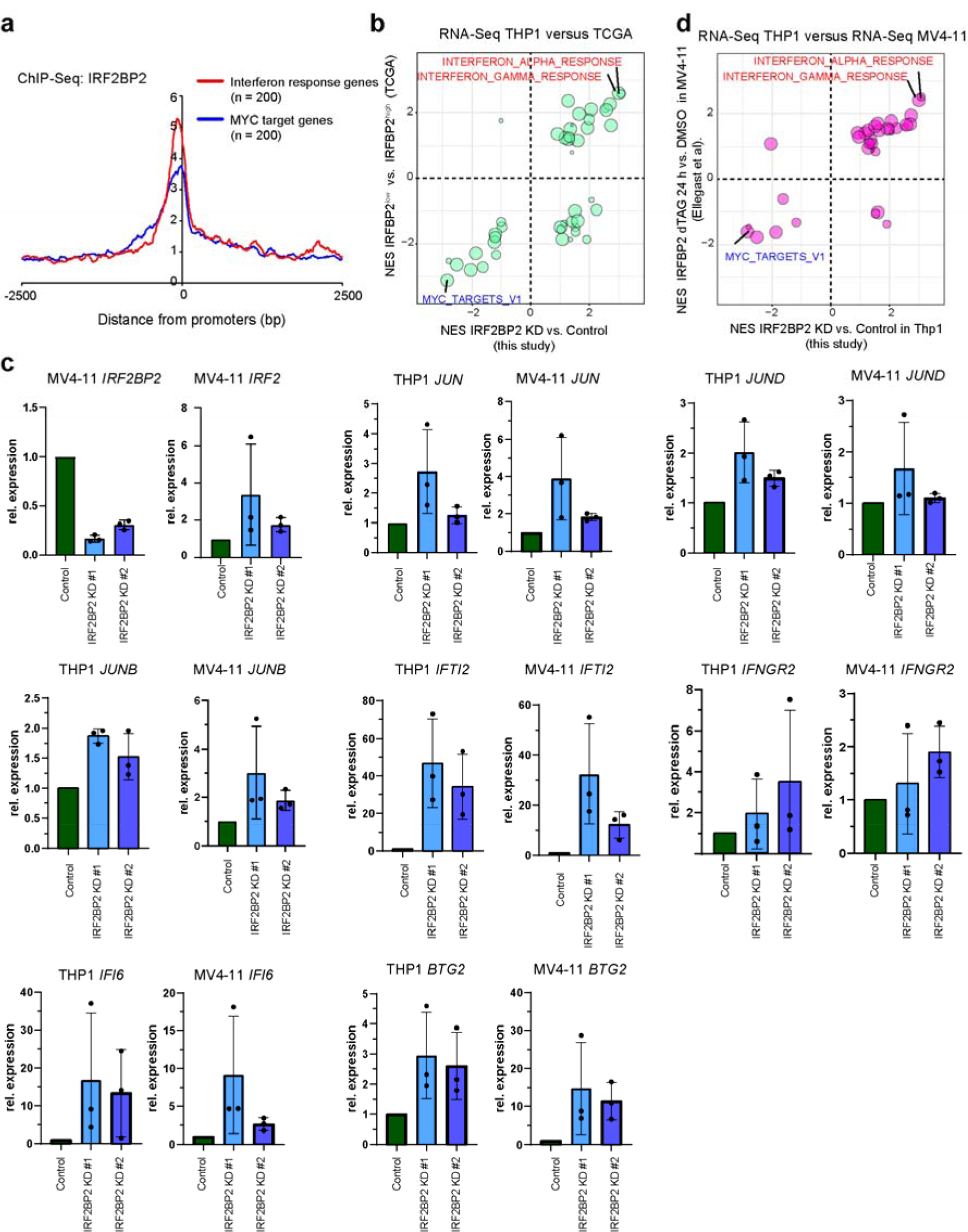
Effects of IRF2BP2 deletion is consistent in multiple cell lines. a) ChIP-Seq profile of IRF2BP2 at MYC target genes and interferon response genes (based on the GSEA hallmark dataset). b) Bubble plot comparing the results from GSEA obtained from our RNA-Seq data (IRF2BP2 knockdown versus Control) and analyzing TCGA samples (IRF2BP2^low^ versus IRF2BP2^high^). c) RT-qPCR analysis of inflammatory genes in THP1 and MV4-11 cells. Data represent the mean +/− s.d. of three biological replicates. d) Bubble plot comparing the results from GSEA obtained from our RNA-Seq data (IRF2BP2 knockdown versus control in THP1 cells) and RNA-Seq data from Ellegast et al. (IRF2BP2 dTAG versus DMSO in MV4-11 cells for 24 h).

**Supplementary Fig. 7.**
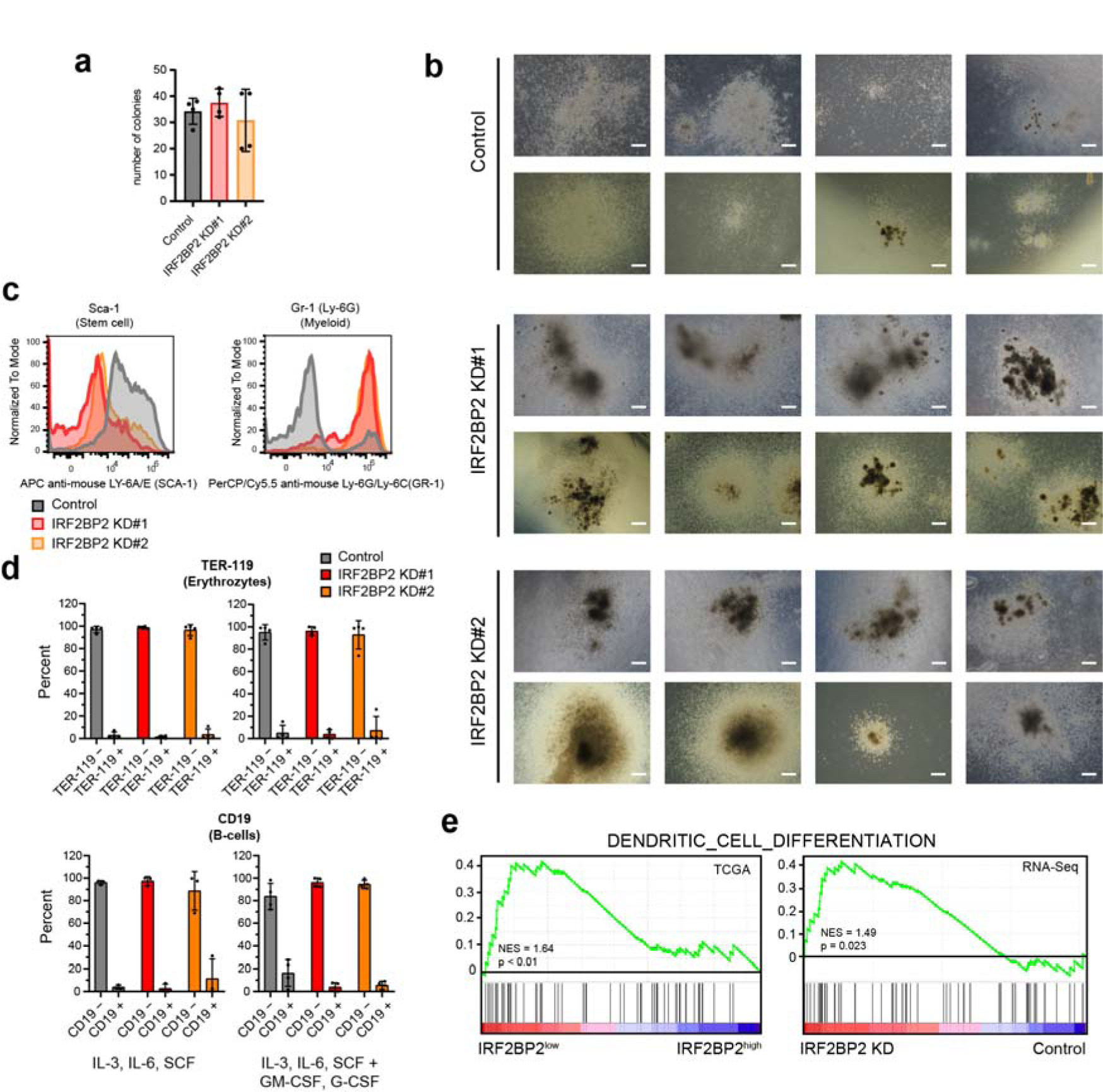
Additional data from differentiation of myeloid stem and progenitor cells. a) Quantification of colonie numbers from methylcellullose assay. Data represent the mean +/− s.d. of at two biological replicates, with two technical replicates, respectively. b) Additional examples of colonies obtained upon differentiation in methylcellulose. Scale = 500 µm. Related to figure 7b. c) Exemplary histograms of IL-3, IL-6, SCF, GM-CSF and G-CSF differentiated myeloid progenitor cells, stained with Sca-1 and Gr-1. d) FACS quantification of surface markers for B-cells (CD19) and erythrocytes (TER-119) after 10 days of differentiation of Lin-negative bone marrow cells. Data represent the mean +/− s.d. of at least three biological replicates. e) GSEA of TCGA and our own RNA-Seq data regarding the “Dendritic Cell Differentiation” pathway.

**Supplementary Table S1:** Overview of used antibodies

| Antibody | Source | Catalog number |
| --- | --- | --- |
| IRF2BP2 | Sigma Aldrich | HPA027815 |
| IRF2BP1 | Proteintech | 13698-1-AP |
| IRF2BP1 | Sigma | HPA049480 |
| ATF7 | Bethyl Laboratories | A302-431A |
| JDP2 (3C1) | Santa Cruz | sc-517133 |
| FLAG (mouse) | Sigma Aldrich | F3165 |
| SP-1 (rabbit) | Self-made | 53 |
| H3 | Abcam | ab1791 |
| HA (rat) | Merck | 11867423 |
| Beta Tubulin (KMX-1) | Millipore | MAB3408 |
| Beta Actin | Santa Cruz | sc-47778 |
| KDM1A | Abcam | AB17721 |
| Mouse(sheep)HRP | GE Healthcare | NA931 |
| Rabbit (donkey)HRP | GE Healthcare | NA934 |
| Rabbit (goat) Alexa Fluor 488 | Thermo Fisher Scientific Inc. | A-11008 |
| Rat (goat) HRP | GE Healthcare | NA935 |
| Rat (donkey) Alexa Fluor 488 | Thermo Fisher Scientific Inc. | A-21208 |
| Rat (goat) Alexa Fluor 546 | Thermo Fisher Scientific Inc. | A-11081 |
| APC anti-mouse LY-6A/E (SCA-1) | Biozol | BLD-122512 |
| PerCP/Cy5.5 anti-mouse CD117 (c-Kit) | Biozol | BLD-105824 |
| APC anti-mouse/human CD11b | Biozol | BLD-101212 |
| PerCP/Cy5.5 anti-mouse Ly-6G /Ly-6C(GR-1) | Biozol | BLD-108428 |
| APC anti-mouse /human CD19 | Biozol | BLD-115512 |
| PerCP/Cy5.5 anti-mouse TER-119/Erythroid Cells | Biozol | BLD-116228 |
| APC Rat IgG2a kappa Isotype Ctr, Clone RTK2758 | Biozol | BLD-400512 |
| PerCP/Cy5.5 Rat IGG2b, kappa Isotype Ctr, Clone: RTK4530 | Biozol | BLD-400632 |
| APC Rat IgG2b kappa Isotype Ctr, Clone RTK4530 | Biozol | BLD-400612 |

**Supplementary Table S2:**
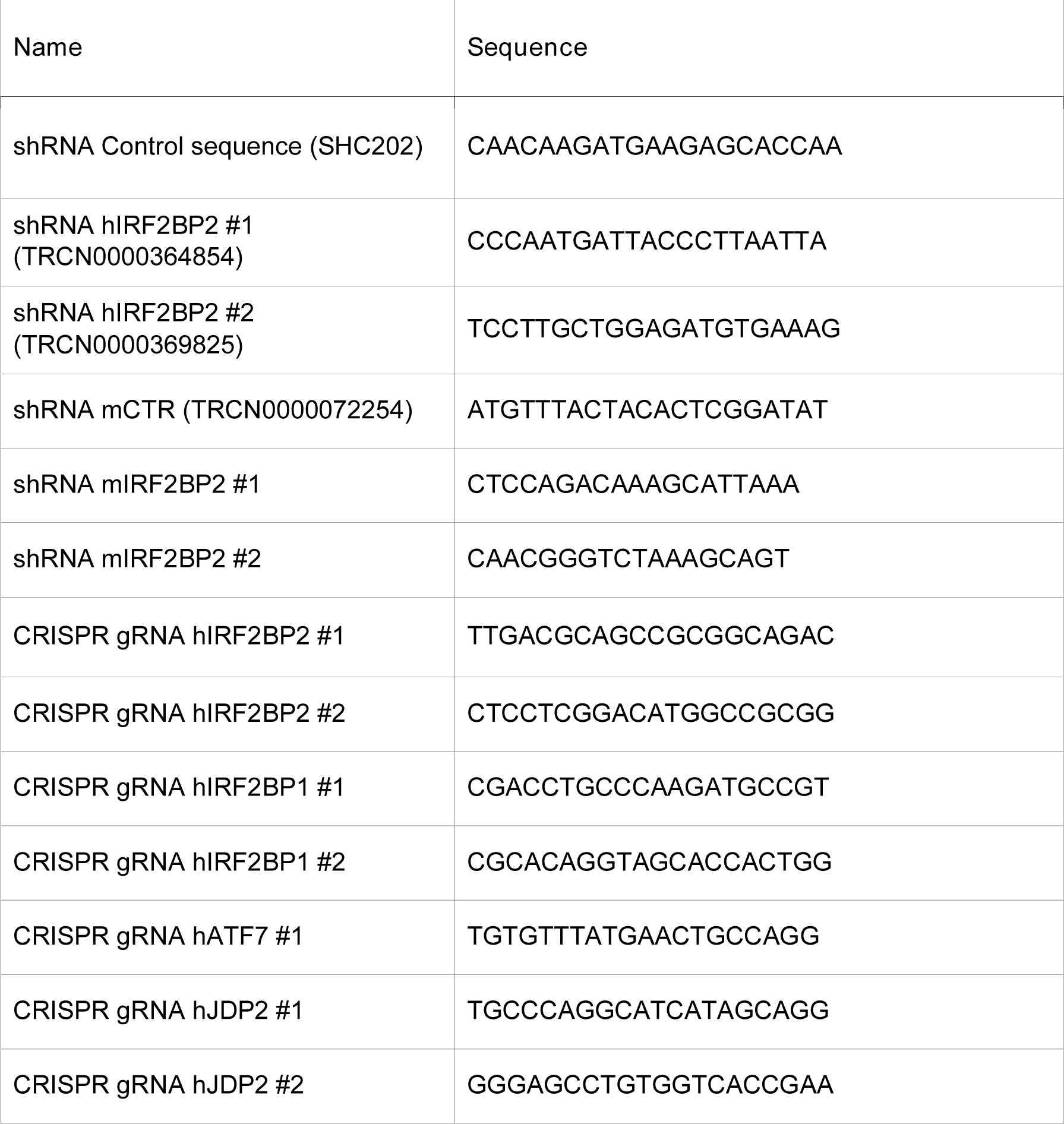
Sequences for shRNA-mediated knockdown and CRISPR-mediated knockouts

**Supplementary Table S3:**
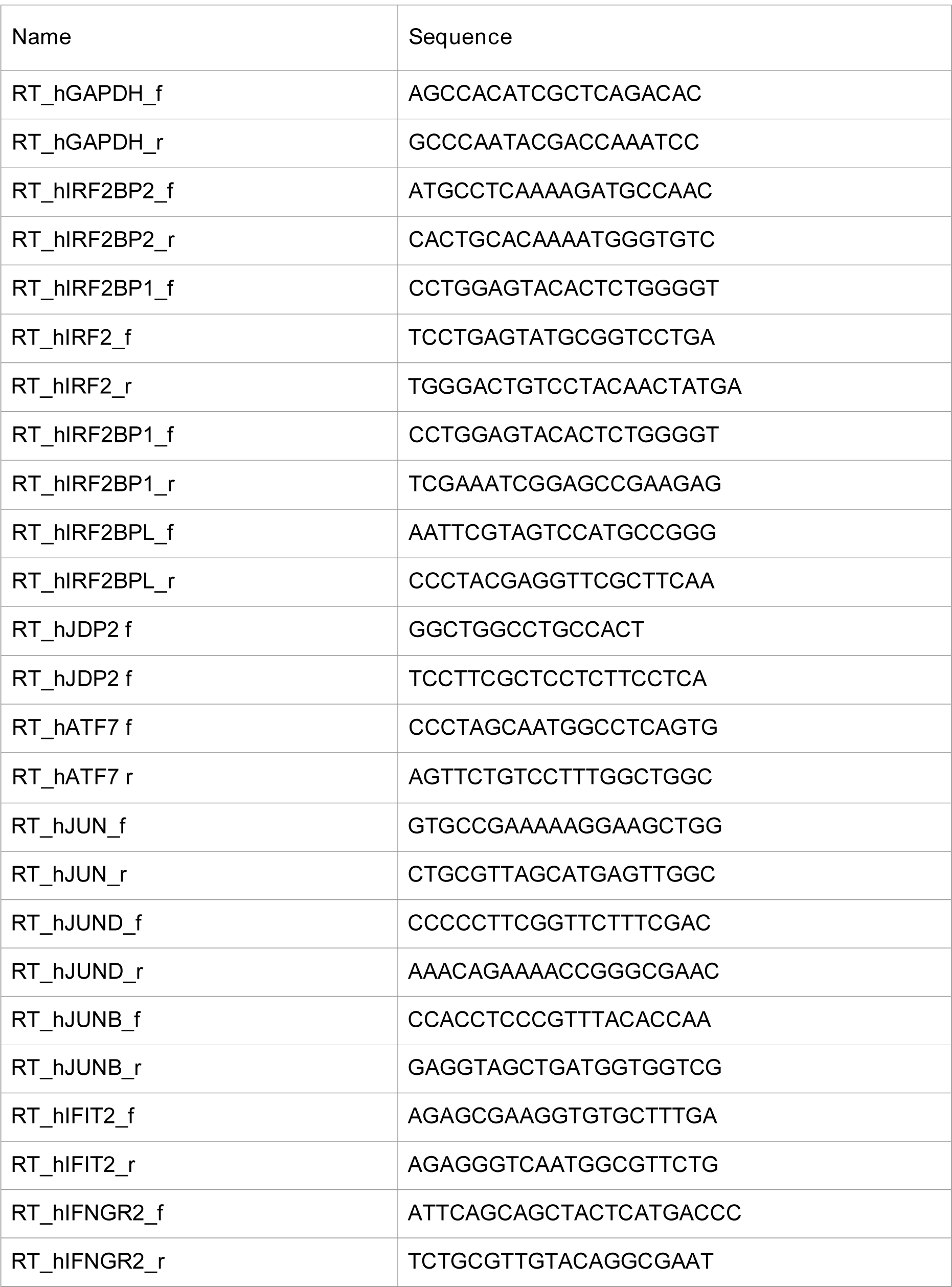
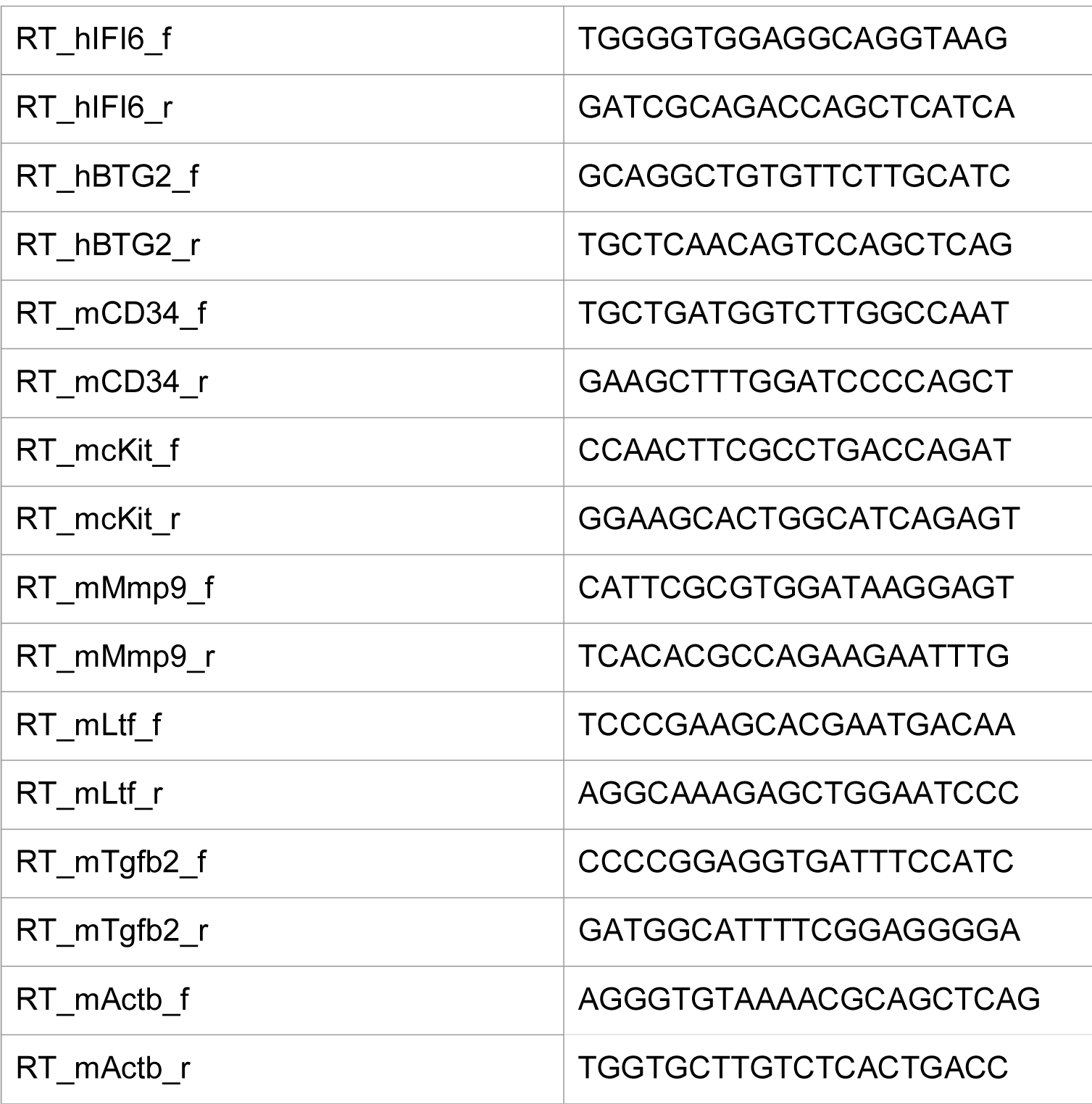
Primers for RT-qPCR

**Supplementary Table S4:**
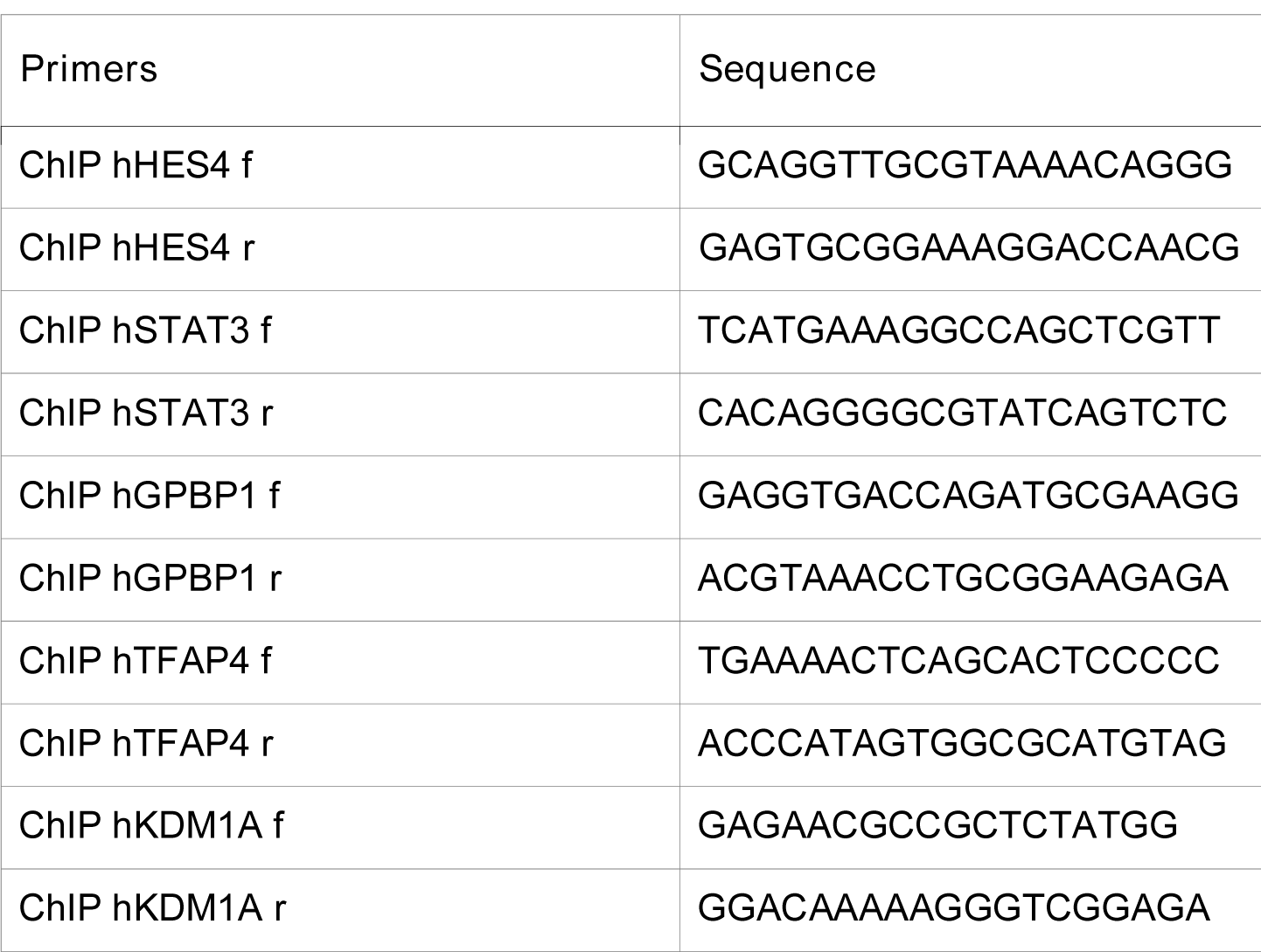
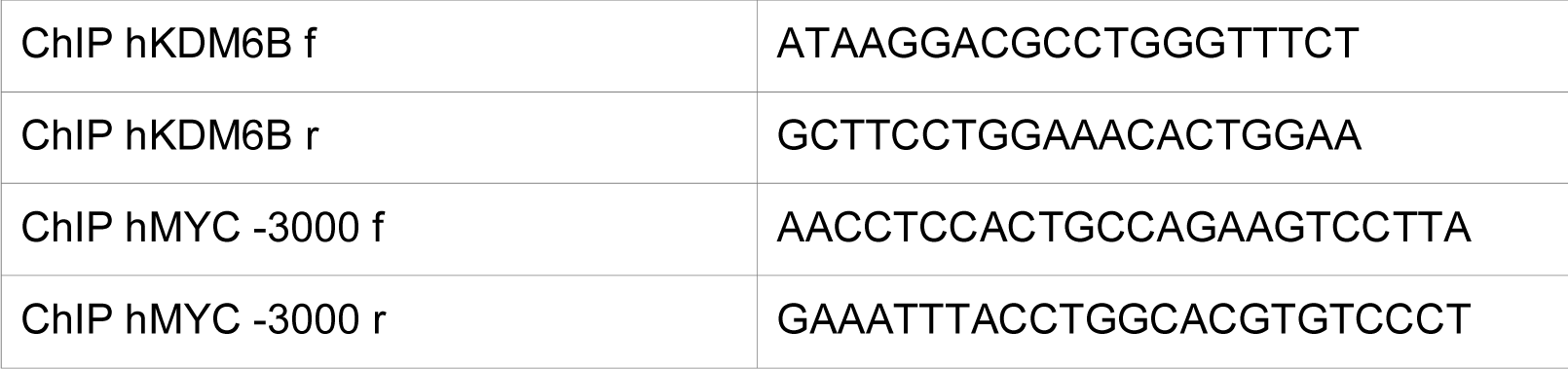
Primers for ChIP-qPCR

**Supplementary Table S5:**
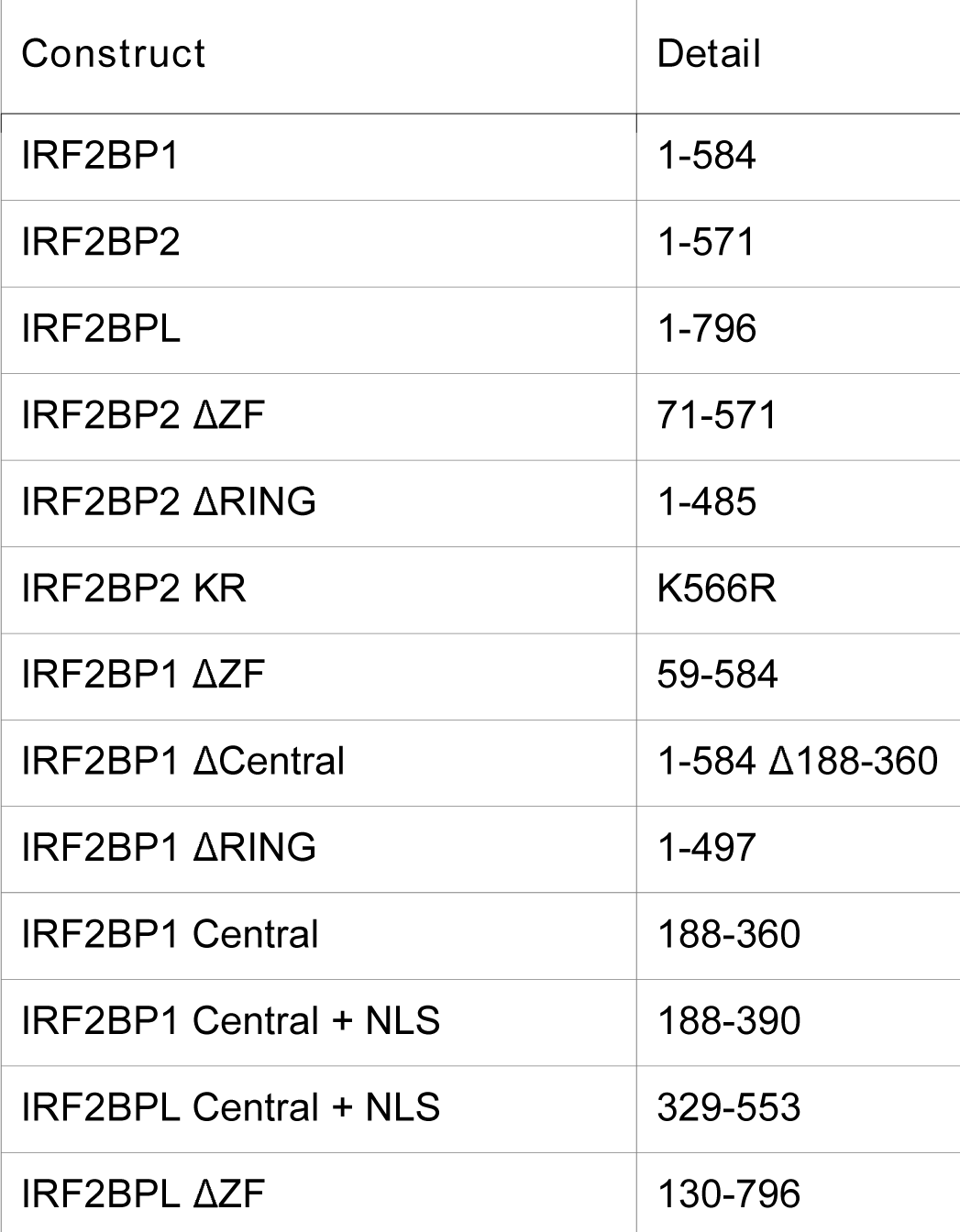
Overview of used constructs

| Construct | Detail |
| --- | --- |
| IRF2BP1 | 1-584 |
| IRF2BP2 | 1-571 |
| IRF2BPL | 1-796 |
| IRF2BP2 ΔZF | 71-571 |
| IRF2BP2 ΔRING | 1-485 |
| IRF2BP2 KR | K566R |
| IRF2BP1 ΔZF | 59-584 |
| IRF2BP1 ΔCentral | 1-584 Δ188-360 |
| IRF2BP1 ΔRING | 1-497 |
| IRF2BP1 Central | 188-360 |
| IRF2BP1 Central + NLS | 188-390 |
| IRF2BPL Central + NLS | 329-553 |
| IRF2BPL ΔZF | 130-796 |

